# Scarless engineering of the *Drosophila* genome near any site-specific integration site

**DOI:** 10.1101/2020.08.13.249656

**Authors:** Siqian Feng, Shan Lu, Wesley B. Grueber, Richard S. Mann

## Abstract

We describe a simple and efficient technique that allows scarless engineering of *Drosophila* genomic sequences near any landing site containing an inverted attP cassette, such as a *MiMIC* insertion. This 2-step method combines phiC31 integrase mediated site-specific integration and homing nuclease-mediated resolution of local duplications, efficiently converting the original landing site allele to modified alleles that only have the desired change(s). Dominant markers incorporated into this method allow correct individual flies to be efficiently identified at each step. In principle, single attP sites and FRT sites are also valid landing sites. Given the large and increasing number of landing site lines available in the fly community, this method provides an easy and fast way to efficiently edit the majority of the *Drosophila* genome in a scarless manner. This technique should also be applicable to other species.

## Introduction

Reverse genetics is a powerful tool to study the functions of genes and proteins. To answer many important biological questions, it is necessary to make precise genomic changes at the base pair resolution, preferably in a scarless manner, such that the final alleles only have the desired mutation(s). It is therefore important to have simple and efficient techniques for scarless genome engineering.

The fruit fly *Drosophila melanogaster* is well known for its superior genetic tool kit. There have been many efforts to precisely engineer the *Drosophila* genome. The first successful attempt used so-called ends-in targeting by homologous recombination to generate a local duplication, followed by homing nuclease-mediated resolution of the duplication (Rong et al., 2002). The final mutant alleles are scarless, but because of the low efficiency of ends-in targeting, large scale screening of thousands of vials is necessary to identify the successful targeting events. A variant technique called SIRT (Site-specific Integrase mediated Repeated Targeting) is suitable for generating multiple different mutant alleles of the same locus (Gao et al., 2008). It involves an initial labor-intensive ends-in targeting step to insert an attP site near the locus of interest, but all subsequent mutagenesis uses highly efficient phiC31 integrase mediated site-specific integration and homing nuclease-mediated resolution of the duplication. The final alleles generated by SIRT still have an attR scar.

RMCE (recombinase mediated cassette exchange) (Bateman et al., 2006) based techniques represent a different strategy (Delker et al., 2019). In these approaches, the wild type locus is first replaced by an inverted attP cassette, two attP sites in the opposite orientation flanking a dominant marker. This is usually achieved by homologous recombination induced by cutting with a custom endonuclease such as ZFN, TALENs, or CRISPR. Next, phiC31 integrase mediated RMCE is used to replace the dominant marker with a mutant version of the genomic sequence. RMCE based techniques are relatively straightforward to perform and highly efficient, but the final alleles have two attR scars flanking the modifications.

Most recently, the CRISPR revolution has made the precise engineering of the animal genomes significantly easier. In *Drosophila*, to facilitate the identification of correctly engineered individuals, a dominant marker is often inserted into the genome as the wild type sequence is converted into the mutant sequence during CRISPR-mediated homologous recombination (Gratz et al., 2014). The dominant marker can later be removed, but a short scar such as an FRT site or a loxP site, is often left in the genome, although there are ways to remove the dominant marker in a scarless manner (for example with piggyBac, https://flycrispr.org/). In principal, scarless mutant alleles can also be directly generated by CRISPR-mediated homologous recombination. However, since most custom mutant alleles do not have easily observable phenotypes, individuals bearing the desired mutations must be identified by laborious molecular screening, and when the desired mutation only affects a few base pairs, or even a single base pair, PCR primers may not be able to distinguish the wild type and mutant sequences. In addition, a common challenge with CRISPR based experiments is that the efficiency of the selected gRNA(s) is difficult to predict, and the rate of unsuccessful CRISPR attempts is not trivial (Kanca et al., 2019). Common strategies to increase gRNA efficiency are to test them in cell culture before injecting flies, or to generate gRNA expressing transgenic flies (Port et al., 2015), both of which require additional time and effort.

Here, we report a new approach that combines phiC31 integrase mediated RMCE and homing nuclease mediated resolution of local duplications to scarlessly engineer the *Drosophila* genomic sequences near any landing site with an inverted attP cassette. In this method, first a properly marked mutant DNA fragment is integrated into the selected landing site via RMCE. This creates local duplications on both sides of the integration sites, which are then resolved in a single step by homing nuclease-induced homologous recombination between the duplications, resulting in scarless mutant alleles. Previously, there have been some attempts to combine these two procedures for genome engineering. For example, Zolotarev *et. al.* resolved one side of an RMCE allele in a scarless manner, while the other side still had a scar (Zolotarev et al., 2019). Vilain *et. al.* resolved the two sides one at a time to make scarless alleles, but this method did not include any visible marker, and relied entirely on molecular methods to identify the desired mutation (Vilain et al., 2014). To our knowledge, there have been no reports describing the simultaneous resolution of both sides after RMCE, which significantly shortens the time required to generate the final scarless allele. Once an RMCE line has been generated, our method takes less than two months to obtain a final scarless allele.

Because of the large number of fly lines with inverted attP cassettes, a significant portion of the *Drosophila* genome is accessible with this technique. There are about 17,500 *MiMIC* insertion lines (Lee et al., 2018, Nagarkar-Jaiswal et al., 2015, Venken et al., 2011), and 7441 have been mapped. The mapped *MiMIC* insertions allow approximately half of the euchromatic *Drosophila* genome to be efficiently engineered with this method (see Discussion). The fact that single attP sites and FRT sites are also potential landing sites further expands the accessible portion of the fly genome. phiC31 integrase mediated site-specific recombination, such as RMCE, has been proven to be robust and efficient, and does not have the risk associated with CRISPR gRNA selection. We show that this technique can be used to efficiently make precise protein coding mutations as well as large insertions and deletions. This technique requires no laborious screening and efficiently generates the desired scarless alleles in a short period of time.

## Results

### Overall strategy

Figure 1 illustrates the overall 2-step strategy of this method, exemplified using a landing site containing an inverted attP cassette, such as a *MiMIC* (Nagarkar-Jaiswal et al., 2015, Venken et al., 2011) or CRIMIC insertion (Lee et al., 2018). First, a fragment from the targeting plasmid is integrated into the selected landing site via RMCE. This fragment contains the desired mutation(s), and is flanked by dominant markers and homing nucleases sites. Second, the homing nucleases are expressed, and the resulting dsDNA breaks induce homologous recombination between the integrated mutant sequence and the original genomic sequences, thus resolving the locus to scarless mutant alleles. Alternatively, the two sides can be resolved sequentially (Figure 1-figure supplement 1). In each step, desired individuals are identified by the presence or absence of dominant markers, which greatly simplifies the screening process. Importantly, the final alleles only have the desired mutation(s), with no additional modifications.

**Figure 1.**
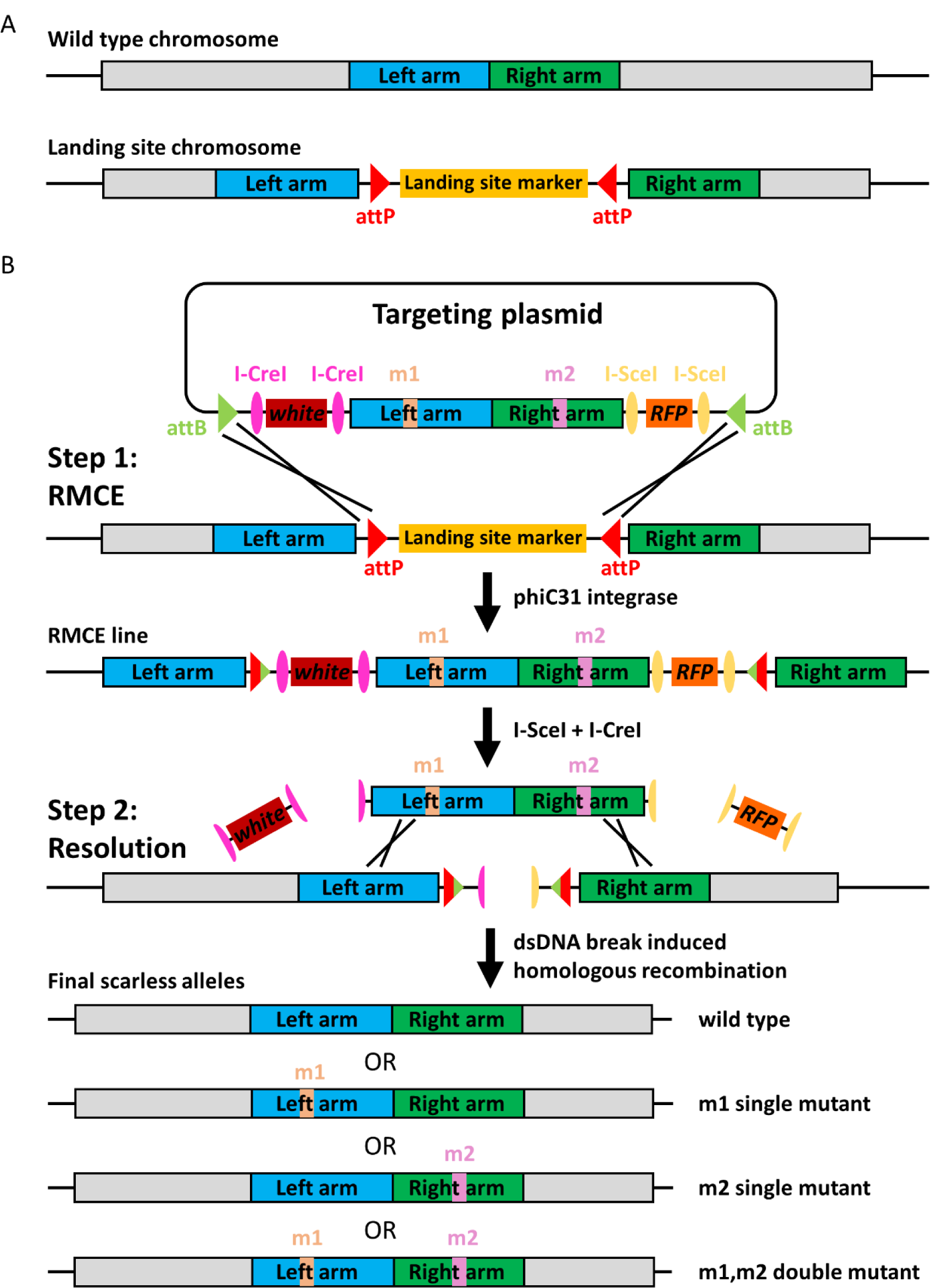
Overall strategy of the genome editing method. **A.** Schematics showing the wild type chromosome and the landing site chromosome. **B.** Schematic that details the 2-step genome editing strategy. In step 1, a properly marked DNA fragment with the desired mutation(s) is integrated near the locus of interest. In step 2, homologous recombination induced by homing nuclease generated dsDNA breaks resolves the local duplications, and generates the final scarless mutant alleles.

### Test of principle: engineering of the *Antp* locus by sequential resolution

The *Hox* gene *Antennapedia (Antp)* was selected for an initial test of this technique. There is a *MiMIC* insertion (*Antp*^*MI02272*^) in the intron between the first coding exon and the small second coding exon, where the so-called W-motif is located (Figure 2A) (Merabet and Mann, 2016). The W-motif, also called the hexapeptide, is a protein-protein interaction motif present in nearly all Hox proteins that mediates the interaction between Hox proteins and their shared cofactor, the TALE family homeodomain protein (Extradenticle (Exd) in *Drosophila* and Pbx in vertebrates) (Mann et al., 2009). Although the functions of the W-motif have been extensively studied, most *in vivo* experiments rely on ectopic expression of mutant Hox proteins (Merabet and Mann, 2016). Therefore, this motif represents an interesting target for *in vivo* genome engineering.

**Figure 2.**
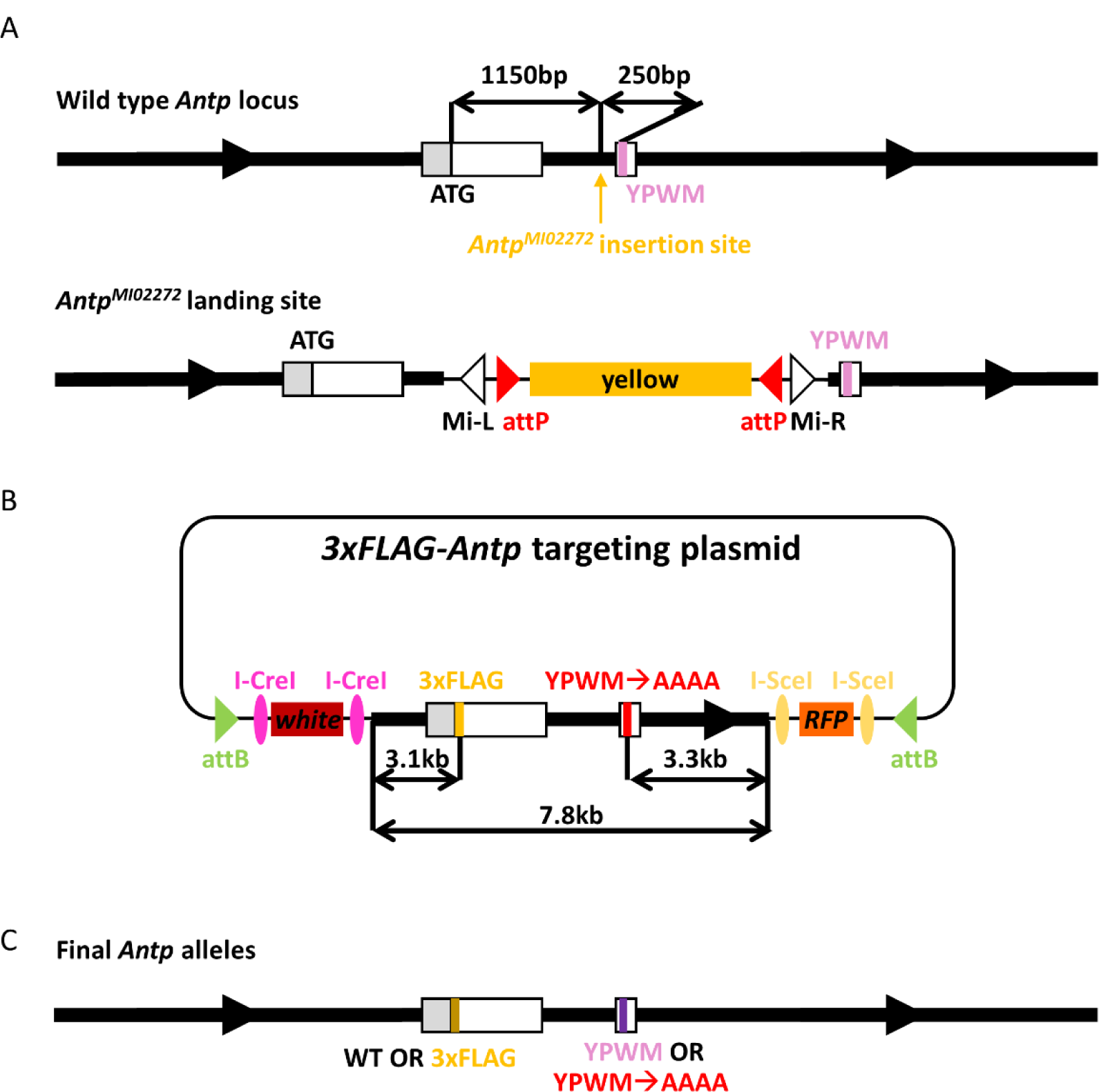
Scarless engineering of the *Antp* locus. **A.** The wild type *Antp* locus and the *AntpMI02272 MiMIC* landing site. The thick black lines denote introns and the arrows indicate the direction of transcription. The white boxes are coding exons, and the gray box shows part of the 5’ UTR. The ATG start codon and the sequence encoding the W-motif, as well as their distances to the *MiMIC* insertion site, are indicated. **B.** The *Antp* targeting plasmid. The total length of the integrated fragment, as well as the relative positions of the two desired mutations, are shown. **C.** The final scarless alleles. The schematics in this figure are not drawn to scale.

The *Antp*^*MI02272*^ *MiMIC* insertion is 1150 bp downstream of the ATG start codon, and 250 bp upstream of the W-motif codons (Figure 2A). Our goals were to insert a short 3xFLAG tag at the N terminus of the Antp protein and to mutate the W-motif from YPWM to AAAA (4 alanines) (Figure 2C). To generate the targeting plasmid, a 7.8 kb genomic fragment flanking the *MiMIC* insertion site, which had the N terminal 3xFLAG tag and the YPWM->AAAA mutation, was cloned into the optimized targeting vector, which contains the required markers and homing endonuclease sites (Figure 2B and Figure 1-figure supplement 2). This targeting plasmid was injected into the F1 embryos of the cross between the *MiMIC* males and females from the *vas-int(X)* line, which specifically expresses the phiC31 integrase in the germline (Bischof et al., 2007) (see Methods for details).

Successful RMCE events were identified by the presence of both *mini-white* and *3xP3-RFP* markers, as well as the simultaneous loss of the *yellow* marker present in the original *MiMIC* insertion. PCR was performed to determine the orientation of the RMCE lines. The entire targeting plasmid was about 20 kb in size, and a 17 kb fragment was integrated into the genome through RMCE. Despite the large size, multiple independent RMCE lines with the correct orientation were readily obtained. Due to the presence of insulators with repetitive sequences (see Figure 1-figure supplement 2 and Materials and Methods), Southern blot analysis was performed to ensure there were no unwanted rearrangements (Figure 2-figure supplement 1). One fully verified RMCE line was selected for the next resolution step.

Prior to testing the simultaneous resolution of both sides, we first tested the sequential resolution of each side (Figure 1-figure supplement 1). The right side was resolved first by expressing the homing endonuclease I-SceI (Figure 3A), which has an 18 bp recognition site that is not present in the *Drosophila* genome (Bellaiche et al., 1999). The *hs-I-SceI* flies were crossed to the RMCE flies (cross I) (Figure 3A), and their F1 embryos/larvae were heat-shocked at 37°C for 1 hour to induce I-SceI expression. 100 F1 adult males were then individually crossed to a balancer stock (cross II) (Figure 3A). Every fertile cross II produced at least one male progeny that had lost the *3xP3-RFP* marker, suggesting a high efficiency. To ensure all resolved lines were independent, only one male that lost *3xP3-RFP* from each individual cross II was selected to generate a stock (cross III) (Figure 3A).

**Figure 3.**
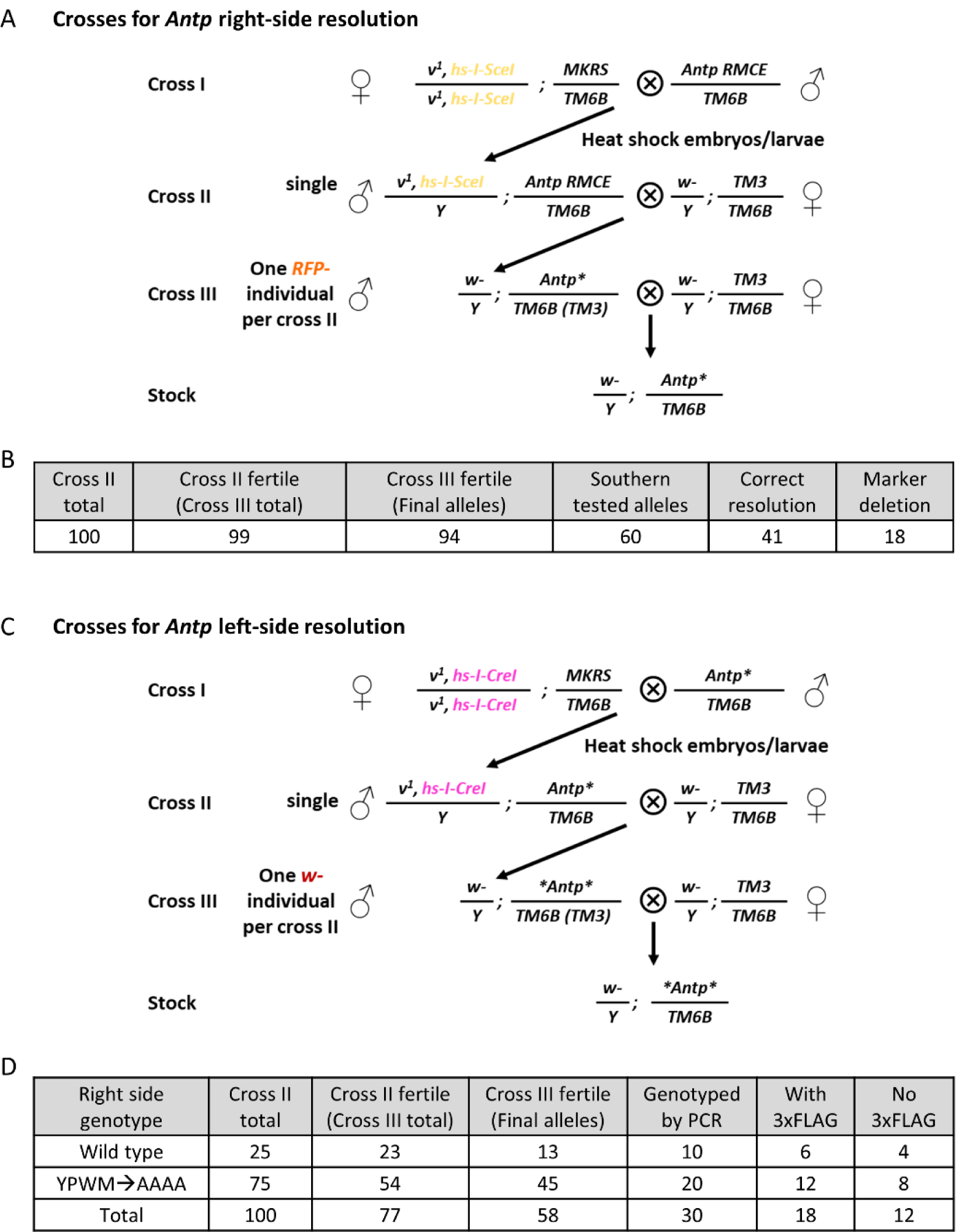
Sequential resolution of the *3xFLAG-Antp* RMCE allele. **A.** The crosses for I-SceI mediated right-side resolution of the *3xFLAG-Antp* RMCE allele. **B.** The results of I-SceI mediated right-side resolution. Of the 94 independent final alleles, 60 were randomly selected for Southern blot analysis. **C.** The crosses for I-CreI mediated left-side resolution. **D.** The results of I-CreI mediated left-side resolution. 30 out of 58 final alleles were genotyped by PCR.

In total, 94 independent right side-resolved lines were obtained, and 60 lines were randomly selected for Southern blot analysis. 41/60 lines showed the expected pattern of a successful resolution. The patterns of 18 of the other 19 lines were consistent with a marker deletion event induced by two double stranded DNA breaks flanking the *3xP3-RFP* marker (Figure 3B and Figure 3-figure supplement 1). Out of the 41 successfully resolved lines, 38 had the YPWM->AAAA mutation, and 3 were wild type, as determined by genotype-specific PCR. The DNA sequence encoding the YPWM->AAAA mutation is 250 bp from the *MiMIC* insertion site and about 3300 bp from the end of the right homologous arm (Figure 2A and 2B). Remarkably, the 3:38 observed wild type to mutant ratio is very close to expectations (250:3300 ≈ 3:40) if recombination is evenly distributed across the homology arm. Finally, we sequenced 2 wild type and 2 mutant alleles between the *MiMIC* insertion site and the end of the right homologous arm, and confirmed that no unwanted mutations were introduced.

Next, from fully verified right-side resolved lines, one wild type line and one mutant line were selected for left side resolution by I-CreI. The overall resolution strategy and crosses were essentially the same as for the right-side resolution (Figure 3C). I-CreI has endogenous target sites in the 28S rRNA gene in the heterochromatic regions on X and Y chromosomes (Rong et al., 2002), thus prolonged expression causes high rates of lethality and sterility. To reduce the toxicity of I-CreI, the heat shock was performed for 40 minutes at 37°C. Under such conditions, a mild reduction in fertility was observed in cross IIs, but each fertile cross II still produced at least one *w-* male. A slight reduction in fertility was also seen in cross IIIs (Figure 3D).

10 fully resolved wild type lines and 20 fully resolved mutant lines were selected for further characterization. PCR was used to determine if the 3xFLAG tag was present, and to eliminate any marker deletion lines from further characterization. 60% of the resolved lines had the 3xFLAG tag (Figure 3D), which was close to the expected ratio (∼70%) based on the relative position of the ATG start codon in the integrated fragment (Figure 2A and 2B). 5 independent *3xFLAG-Antp* alleles, 4 independent *Antp(YPWM->AAAA)* alleles, and 7 independent *3xFLAG-Antp(YPWM->AAAA)* alleles were selected for Southern blot verification, and all gave the expected patterns (Figure 3-figure supplement 2).

One noteworthy finding was that marker deletion events were not observed during left side resolution, either by PCR or by Southern blot, in contrast to right side resolution. Detailed inspection of the targeting vector revealed that the two I-SceI sites flanking the right side *3xP3-RFP* marker were in the same orientation, such that the single stranded overhangs generated by I-SceI were compatible, thus facilitating a marker deletion event. In contrast, the two I-CreI sites flanking the left side *mini-white* marker were in the opposite orientation, thus the single stranded overhangs generated by I-CreI were not compatible, disfavoring simple marker deletion events.

### Simultaneous resolution of both sides

Next, we tested the simultaneous resolution of both sides, which would significantly simplify and shorten the entire process (Figure 1B). The overall procedure was similar to left-side or right-side resolution, except that both I-SceI and I-CreI were expressed together. The simultaneous resolution crosses for chromosome III targets are shown in Figure 4A, and those for chromosome II and X targets are shown in Figure 4-figure supplement 1. We tested heat shock at 37°C for 10, 20, 30 and 40 minutes, and found that a 20-minutes heat shock gave the highest rate of productive cross II (data not shown), defined as the fraction of cross IIs that lead to a final stock (see Materials and Methods for more details).

**Figure 4.**
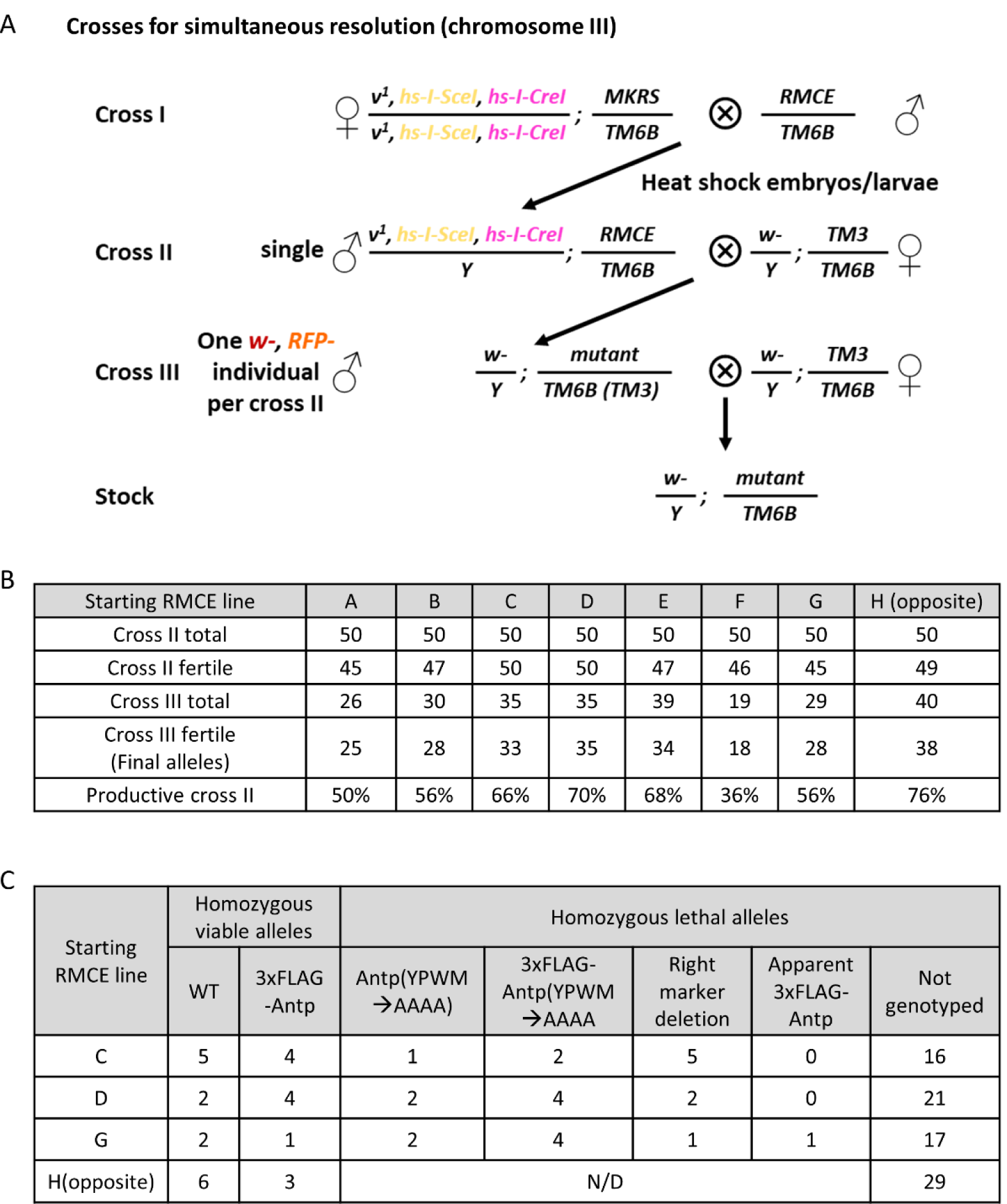
Simultaneous resolution of *3xFLAG-Antp* RMCE alleles. **A.** Crosses for simultaneous resolution of RMCE alleles on chromosome III. **B.** Results of the simultaneous resolution of 8 independent *3xFLAG-Antp* RMCE alleles. RMCE allele H has the opposite integration orientation. **C.** PCR genotyping results of selected final alleles from 4 starting RMCE alleles. For each genotype, multiple independent alleles were obtained. For RMCE allele H, which has the opposite orientation, only homozygous viable final alleles were genotyped. “N/D” stands for “not determined”.

To gain a better measure of the efficiency and robustness of this method, 8 different verified RMCE lines were subjected to simultaneous resolution (Figure 4B). After a 20-minute heat shock, essentially normal viability and fertility were observed. On the other hand, not all individual cross IIs generated male progeny that lost both the *mini-white* and the *3xP3-RFP* markers (Figure 4B); as expected, we frequently observed cross II progeny that lost either *mini-white* or *3xP3-RFP*, but not both. Nevertheless, except for one RMCE line (line F), the rate of productive cross II ranged from 50% to 70%, confirming the high efficiency of simultaneous resolution (Figure 4B).

We selected the final alleles resolved from 3 different RMCE lines for further characterization. PCR was first used to genotype the selected alleles. Because Antp’s W-motif motif is expected to be necessary for viability, only the presence of the *3xFLAG* tag was examined for all homozygous viable final alleles. For selected homozygous lethal alleles, the presence of the *3xFLAG* tag, the *YPWM->AAAA* mutation, as well as the potential right-side marker deletion were tested (Figure 4C). We detected some right-side marker deletion events, as expected. One homozygous lethal allele had the apparent genotype of *3xFLAG-Antp+*. Presumably, an unwanted mutation occurred during resolution, which caused the observed homozygous lethality. Southern blot was performed on 15 genotyped alleles, and 14 gave the expected patterns (Figure 4-figure supplement 2), confirming the high accuracy of this technique. Finally, all 14 Southern blot-verified lines were confirmed by sequencing, and contained no unwanted mutations.

### RMCE lines with opposite integration orientation can be resolved efficiently

The observation that simultaneous resolution works well raised the possibility that successful resolutions can be obtained even if the original RMCE line was in the opposite orientation, where the duplicated arms are not adjacent to their endogenous homologous sequences. We considered this possibility because when both I-SceI and I-CreI are expressed, the entire integrated fragment is liberated from the chromosome and, in principle, could pair with homologous sequences regardless of the initial orientation. We tested this with an RMCE line in the opposite orientation (Figure 4B). Indeed, this line showed a resolution efficiency that was among the highest of all 8 tested RMCE lines.

To confirm the accuracy of the final alleles, we further characterized all 9 homozygous viable alleles generated from this particular RMCE line. Of these 9 alleles, 3 had the *3xFLAG-Antp* genotype, while the other 6 were untagged (Figure 4C). We selected 2 of the 3 *3xFLAG-Antp* alleles for further verification by Southern blotting, and both gave the expected patterns (Figure 4-figure supplement 2). The sequences of these 2 alleles confirmed that there were no additional mutations.

### Precise editing of *Ubx*, another *Hox* gene

To test the generality of this method, we engineered another Hox gene, *Ultrabithorax (Ubx)*. We chose to mutate the canonical W-motif, the YPWM motif, and insert an N terminal 3xFLAG tag (Figure 5A and 5C). As no landing site insertions were available, we first inserted an inverted attP cassette marked with *ubi-DsRed* (Handler and Harrell, 2001) into the *Ubx* locus, using a pair of custom TALENs that target the first coding exon of *Ubx* (Figure 5A; see Methods for details).

**Figure 5.**
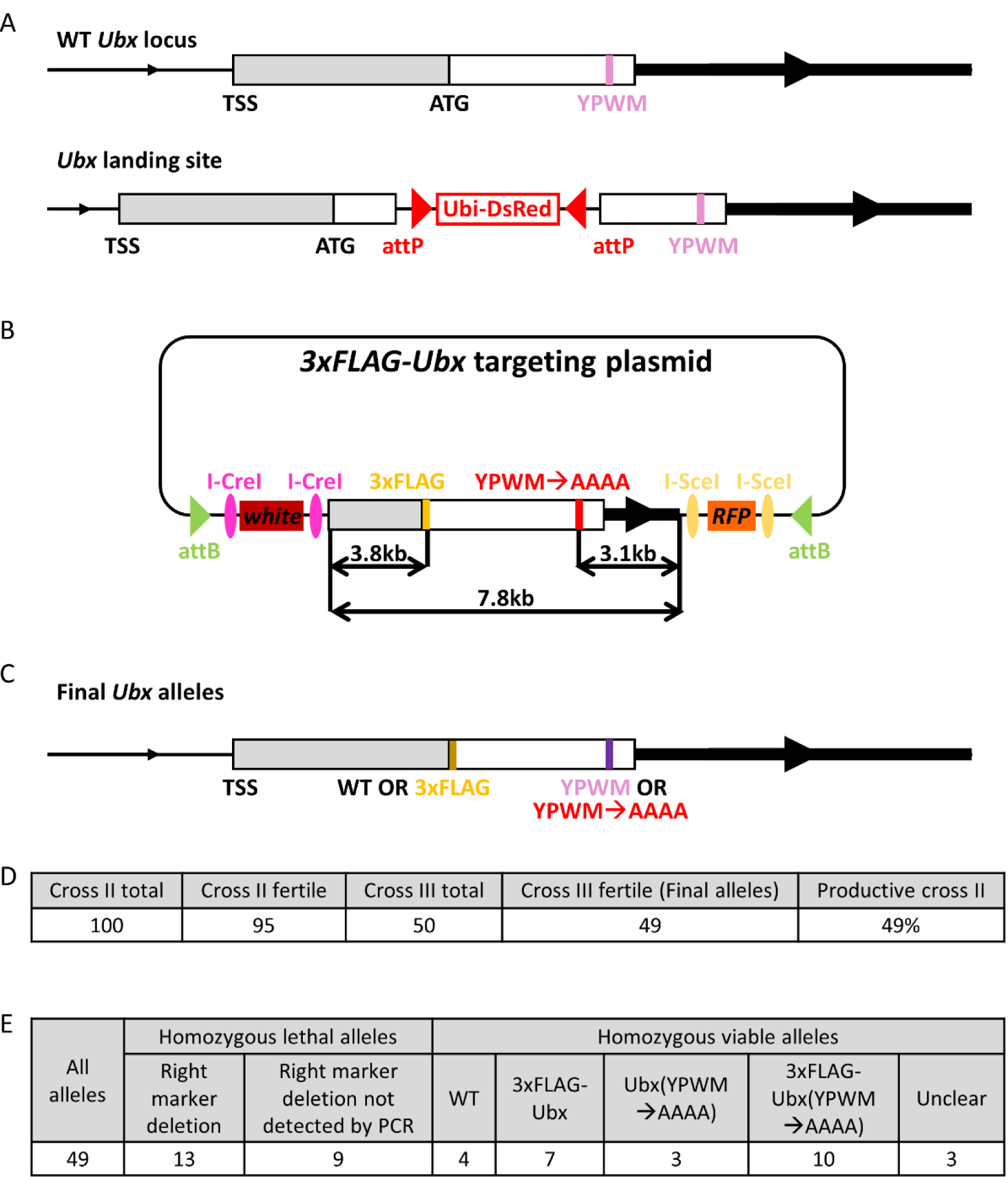
Scarless engineering of the *Ubx* locus. **A.** Schematics of the wild type *Ubx* locus and the *Ubx* landing site allele. **B.** The targeting plasmid used in the scarless engineering of the *Ubx* locus. The desired mutations are shown, as well as their relative positions within the integrated fragment. **C.** The desired final scarless alleles. The schematics in A-C are not drawn to scale. **D.** Results of *Ubx* RMCE allele simultaneous resolution. **E.** Genotyping results of all final *Ubx* alleles. “Unclear” refers to ambiguous genotyping results for 3 homozygous viable alleles.

One fully verified *Ubx* landing site allele was selected as the starting strain for engineering the *Ubx* locus. A *Ubx* targeting plasmid was generated, which contained a 7.8 kb fragment with a 3xFLAG tag at the N terminal end of the *Ubx* ORF and the *YPWM->AAAA* mutation (Figure 5B). This targeting plasmid was injected into the F1 progeny of the *vas-int(X)* females and the *Ubx* landing site males, and multiple independent RMCE lines were obtained and further verified by Southern blot. One fully verified RMCE line was subjected to simultaneous resolution, following the same procedure as for the *Antp* locus. From 100 individual cross IIs, we were able to achieve a success rate of ∼50% (Figure 5D).

Among the 49 alleles obtained, 22 were homozygous lethal, and 27 were homozygous viable (Figure 5E). 13 of the homozygous lethal alleles had the right marker deleted, as shown by PCR. All 4 expected genotypes, wild type, *3xFLAG-Ubx, Ubx(YPWM->AAAA)* and *3xFLAG-Ubx(YPWM->AAAA)*, were identified from the 27homozygous viable alleles (Figure 5E), indicating the YPWM motif of Ubx is not necessary for viability. Although this W-motif is deeply conserved, this result was not unexpected because Ubx has multiple additional Exd-interaction motifs (Lelli et al., 2011, Merabet and Mann, 2016, Merabet et al., 2007), which may be able to partially compensate for the functions of the canonical YPWM motif. Some homozygous lethal alleles did not show a PCR product with primers designed to detect right-side marker deletion events (Figure 5E). These alleles might have undergone imprecise homologous recombination, or the marker deletion might have been accompanied by additional deletions near the dsDNA breaks, such that the primer binding sites were destroyed.

Southern blotting was performed to verify 16 different alleles; of these, 3 showed abnormal patterns (Figure 5-figure supplement 1) and they were discarded. 2 Southern blot verified alleles of each genotype of interest were fully sequenced, and all 6 were correct. The precise engineering of the *Ubx* locus demonstrated again the efficiency and precision of this technique.

### Generating insertions

The above results show that this technique can be used to efficiently mutate small stretches of genomic DNA sequence, or to insert a small fragment into a desired genomic locus. To further test the ability of this technique to generate large custom insertions, we chose to tag the endogenous Antp protein with GFP (Figure 6B). A slightly different *Antp* targeting plasmid, in which the 3x-FLAG tag was replaced with a 750 bp GFP tag (with a flexible linker between the *GFP* and *Antp* ORFs), was generated (Figure 6A), and multiple independent RMCE lines were obtained. One Southern blot-verified RMCE line was then used as the starting line for simultaneous resolution. Because it was unclear if and how the large-sized insertion would affect the resolution success rate, we set up 100 individual cross IIs. Despite the presence of a large insertion, the resolution results had a high rate of success: 70% of cross IIs were productive (Figure 6C).

**Figure 6.**
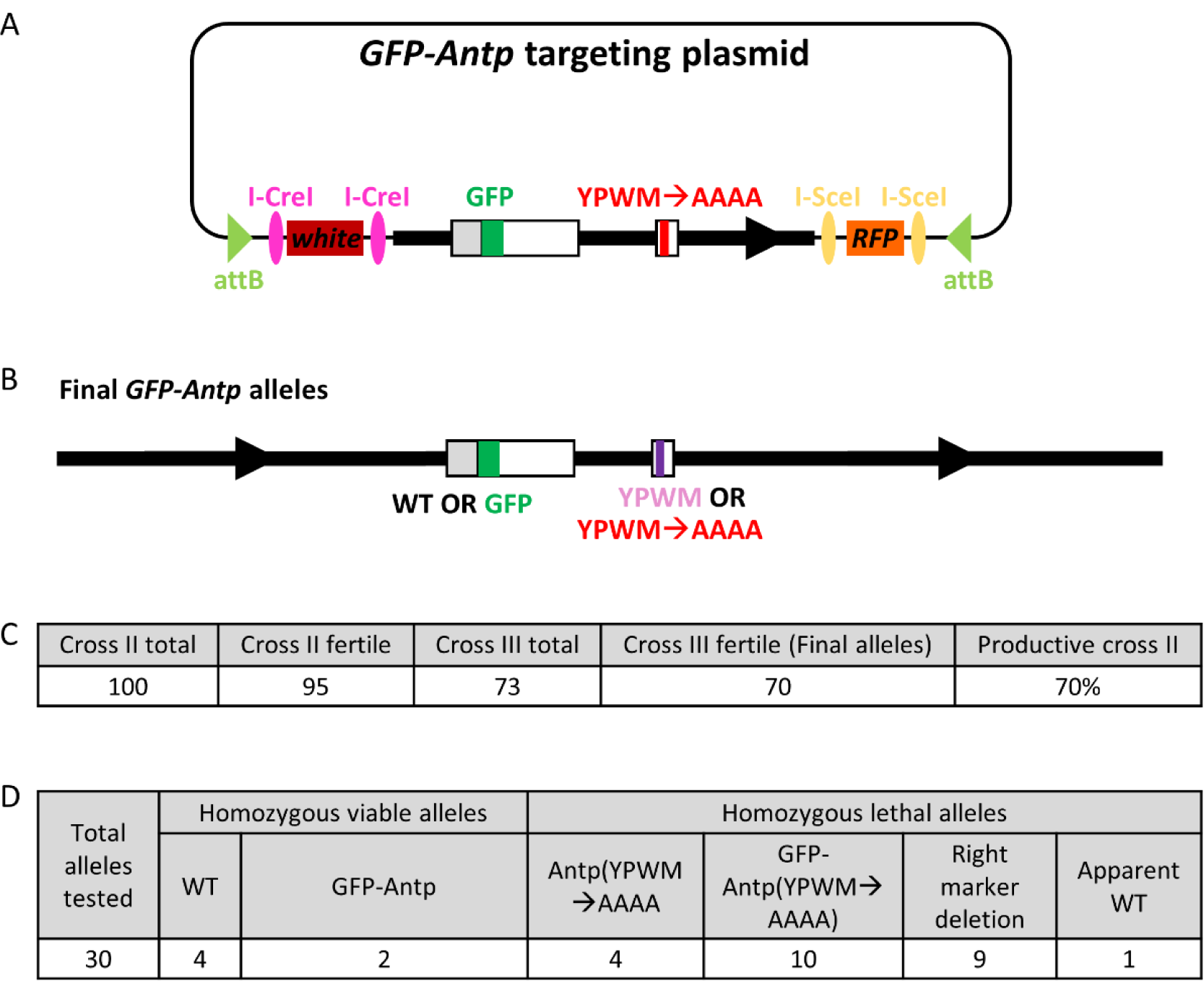
Generating a precise insertion at the *Antp* locus. **A.** The *GFP-Antp* targeting plasmid. The GFP insertion and the YPWM→AAAA mutation are indicated. **B.** Desired final *GFP-Antp* alleles. The schematics in A and B are not drawn to scale. **C.** Results of simultaneous resolution of the selected *GFP-Antp* RMCE allele. **D.** Genotyping results of 30 selected *GFP-Antp* targeting final alleles.

6 of the 70 final alleles were homozygous viable, from which 2 independent *GFP-Antp* alleles were obtained, as determined by PCR, while the other 4 alleles were wild type. Many *Antp(YPWM->AAAA)* and *GFP-Antp(YPWM->AAAA)* alleles were identified by PCR among the homozygous lethal alleles (Figure 6D). Several independent alleles of each genotype were selected for Southern blot verification (Figure 6-figure supplement 1) and all gave the correct patterns. Sequencing results verified that all selected alleles were correct.

### Generating deletions

Finally, to test the ability of this technique to create custom deletions, the 7.5 kb *Gr28b* gene was chosen to be deleted (Figure 7A and 7C). *Gr28b* is a complex gustatory receptor locus that encodes 5 different isoforms, and has been shown to have multiple functions such as thermo-preference and toxin avoidance (Ni et al., 2013, Sang et al., 2019). A *MiMIC* insertion (*MI11240*) about 300 bp away from the right end of the *Gr28b* gene was used as the landing site for the targeted deletion (Figure 7A). A targeting plasmid was generated, which contained a 2 kb fragment to the left of the desired deletion, fused to a 2.3 kb fragment to the right of the desired deletion (Figure 7B). This plasmid was used to inject F1 embryos of the cross between the *vas-int(X)* female and the *MI11240* male, and multiple independent RMCE events were obtained. Unexpectedly, while some RMCE events landed on chromosome II, where the *Gr28b* gene is located, others did not map to this chromosome. The presence of the *MI11240* insertion in the original *MiMIC* stock was verified by PCR before it was used for injection, thus we hypothesized that the original *MiMIC* stock might have a secondary *MiMIC* insertion on a different chromosome. Indeed, genetic crosses indicated the presence of a second *MiMIC* insertion on chromosome IV (data not shown).

**Figure 7.**
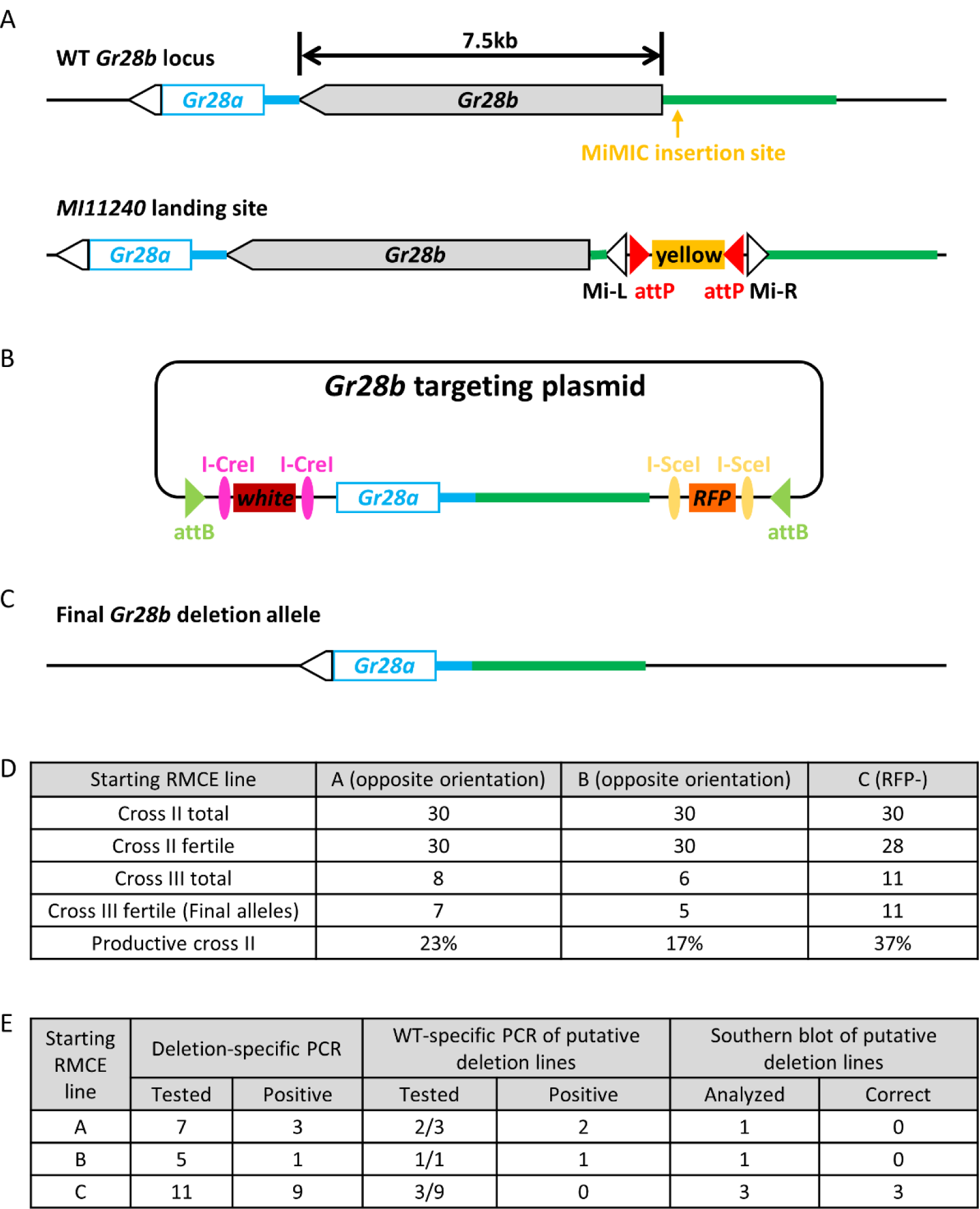
The precise deletion of the 7.5 kb *Gr28b* gene. **A.** Schematics showing the wild type *Gr28b* locus and the chromosome bearing the selected *MiMIC* landing site. **B.** The targeting plasmid containing an integrated fragment with the desired deletion. **C.** Schematic of the desired *Gr28b* deletion allele. The schematics in A-C are not drawn to scale. **D.** Simultaneous resolution results of all 3 RMCE alleles obtained from injection. Two RMCE alleles have the opposite orientation while the third likely underwent spontaneous right-side resolution during RMCE (see text for more details). **E.** The genotyping results of selected *Gr28b* deletion final alleles.

Nevertheless, we obtained 3 independent RMCE events at the desired *MI11240* insertion. Two alleles, A and B, inserted in the opposite orientation, while allele C was *white*+, *yellow*-, but RFP-. The lack of RFP in allele C might be due to spontaneous resolution of the right side induced by the dsDNA breaks generated by the phiC31 integrase during RMCE. Indeed, Southern blot results supported this idea (data not shown), and this allele was essentially equivalent to a right-side resolved allele.

All 3 RMCE alleles were subjected to simultaneous resolution (Figure 4-figure supplement 1A). Multiple independent stocks were obtained from each RMCE line, but compared to the other targeting experiments described above, a notable reduction in efficiency was observed (Figure 7D), likely because the distance between the dsDNA break and the left homologous arm was >7 kb (Figure 7A and 7B) (Gao et al., 2008).

All final alleles were homozygous viable and fertile, and the homozygotes were verified in several steps (Figure 7E). First, the presence of the desired deletion was determined by PCR using primers flanking the deletion. Next, those alleles that generated the correctly sized PCR product were subjected to additional PCRs using two pairs of primers against different regions of the deleted fragment. Several alleles derived from RMCE lines A and B produced positive products for all of these PCRs, suggesting that complex rearrangements occurred during resolution. 3 independent final alleles, all from RMCE line C, passed all PCR tests, and all 3 were further verified by Southern blot (Figure 7-figure supplement 1) and sequencing. Thus, despite a suboptimal initial RMCE step, this technique was able to generate a large 7.5 kb custom deletion.

## Discussion

In the past several decades, research using model organisms has greatly advanced our understanding of biology. Currently, knock-out lines exist for most genes in well studied model organisms, and for future research, precise mutations, such as those affecting only a specific part of a protein, are often necessary to further elucidate the molecular mechanisms underlying various biological processes. Because any scar sequences left in the genome after custom mutagenesis might have unwanted consequences and could confound subsequent analyses, scarless engineering of the genome is often preferred. Here we describe a novel technique that is able to easily and efficiently generate scarless custom mutant alleles in the model organism *Drosophila melanogaster*.

### Advantages of this technique

The advances of CRISPR based techniques have made the engineering of the *Drosophila* genome much easier, but many custom mutant alleles generated with CRISPR still contain sequence scars. Although generating scarless custom mutations in *Drosophila* is feasible, significant effort is required. And regardless of which CRISPR strategy is used, a major uncertainty is that the selected gRNA(s) might be inefficient, or even non-functional. The technique presented here avoids this uncertainty and uses RMCE, a procedure proven to be robust and efficient, to target genomic sequences near the selected landing site.

This technique is simple and fast. The dominant markers ensure the easy identification of desired individuals in each step, and no laborious screening is necessary. If performing the simultaneous resolution, the desired stocks could be obtained in less than two months from the starting RMCE lines.

This technique generates scarless mutant alleles very efficiently. If the desired genomic alterations are not large deletions that necessitate long distances between the dsDNA breaks and the homologous arms during resolution, at least 1/3 of the cross IIs are expected to be productive, and this rate of success is usually much higher, and can even be over 70%. 50 independent cross IIs should assure the successful generation of the desired allele. If multiple combinations of 2 separate modifications at the locus of interest are desired, such as in our *Hox* targeting experiments, increasing the number of cross IIs to 100 should ensure that all desired genotype combinations will be obtained. In fact, this technique is especially suitable for generating multiple combinations of discrete modifications at the locus of interest. Only one injection is performed to obtain an RMCE allele that contains all individual modifications, and the final alleles of all different genotype combinations can be obtained.

This technique is also very robust. Microinjection is a necessary step of essentially any *Drosophila* genome engineering attempt, but microinjection has the potential to result in significant variability. Many factors, such as landing site location, or the presence of a second landing site such as in the case of the *Gr28b* deletion, could lead to suboptimal RMCE injection results. Even if only RMCE lines with opposite orientation, or only lines with spontaneous resolution are obtained and have to be used, the desired alleles can still be generated. The robustness also means that even difficult mutations, such as large deletions, could be generated with this technique, although the efficiencies are expected to be lower compared to simpler modifications.

### This technique can engineer the majority of the *Drosophila* genome

In this study, we did not systematically test how far away from the landing site can be reached and efficiently engineered by this technique. But from previous reports of homing nuclease-mediated resolution of local duplications, we estimate that any sequence within 5 kb from the landing site could be efficiently engineered (Gao et al., 2008, Rong et al., 2002), and sequences as far as 70 kb or even further from the landing site might be engineerable (Wesolowska and Rong, 2013). During resolution, the chromatin could be resolved either to the wild type sequence, or the desired mutant sequence, and the frequency of getting the mutant allele depends on the lengths of the homologous arms, and the distance between the landing site and the locus to be engineered. For loci far from the landing site, it would likely be helpful to increase the length of the homologous arms in the targeting plasmid, such that the arms extend well beyond the locus to be engineered.

There are 17,500 *MiMIC* insertions (Lee et al., 2018, Nagarkar-Jaiswal et al., 2015, Venken et al., 2011) and hundreds of CRIMIC lines (which is steadily increasing) (Lee et al., 2018) that are available to the fly community. Of the 17,500 *MiMIC* insertions, the locations of 7441 are available online (http://flypush.imgen.bcm.tmc.edu/pscreen/downloads.html). 57.5 Mb of the fly genome lies less than 5 kb from a mapped *MiMIC* insertion (see Materials and Methods for the calculation), thus the currently available mapped *MiMIC* insertions provide efficient access to about half of the 117 Mb euchromatic fly genome (Hoskins et al., 2015) by this method. Moreover, these mapped *MiMIC* insertions represent only a subset of all available insertions containing inverted attP cassettes, and insertions with single attP sites, or even FRT sites, are also potential landing sites (see below). Finally, the 5 kb limit for genome modification is also a conservative estimate. Taken together, we estimate that with available landing sites, this method could be used to precisely engineer the majority of the fly genome in a scarless manner. In case there is no suitable landing site near the locus of interest, such as our engineering the *Ubx* locus, a custom landing site can be generated to facilitate scarless genome editing.

### Sequential resolution vs. simultaneous resolution

We have tested two different resolution strategies, sequential resolution and simultaneous resolution. Simultaneous resolution is much faster and can generate the desired alleles from the RMCE lines in less than 2 months. Sequential resolution, on the other hand, takes longer because the one-side resolved alleles must be verified before the second side is resolved. The sequential resolution strategy, however, offers higher efficiency. Except for difficult mutations, essentially over 90% of independent cross IIs were successful, and the failures were only due to sterile male flies. Therefore, when difficult mutations, such as large insertions or deletions, are to be generated, a sequential resolution strategy might be preferred. In fact, to generate the 7.5 kb *Gr28b* gene, all correct deletion alleles were obtained by sequential resolution, except that the first resolution occurred spontaneously during RMCE. When performing sequential resolution, the starting RMCE lines must have the correct orientation, but RMCE lines with the opposite orientation can be used for simultaneous resolution, without an apparent decrease in efficiency.

### Potential extensions of this technique

In this study, only inverted attP cassettes were used as landing sites. It has previously been reported that for homing nuclease mediated resolution of local duplications, resolution efficiency inversely correlated with the distance between homologous arms on chromatin and dsDNA breaks (Gao et al., 2008). Landing sites with inverted attP cassettes are expected to give the highest resolution efficiency, because when RMCE lines from these landing sites are subjected to homing nuclease mediated resolution, only short non-homologous sequences exist between the dsDNA breaks and the homologous arms. However, this does not mean that only inverted attP cassettes can be used as landing sites. Transposon insertions containing a single attP site are also valid landing sites. When using a single attP site as the landing site, the entire targeting plasmid will be integrated into the genome via phiC31 integrase mediated site-specific recombination. The targeting plasmid backbone, as well as extra sequences present in the original attP-containing transposon, will increase the distances between homologous arms on chromatin and the dsDNA breaks. This will likely lead to a decreased resolution efficiency, and aberrant rearrangements might be more frequent (Gao et al., 2008). Nevertheless, given the high efficiency of this technique, we expect that the desired alleles can still be generated.

In addition, flippase (FLP) mediated recombination between FRT sites has been used to integrate plasmids into the *Drosophila* genome in a site-specific manner (Horn and Handler, 2005). In principle, FRT sites could also be used as an initial landing site for this method. However, due to the bidirectional nature of recombination between FRT sites, the plasmid integration efficiency would be expected to be lower than the unidirectional attB-attP integration mediated by phiC31 integrase. Once successful integration events are obtained, the resolution step should work equally well compared to attB-attP integration events. Targeting vectors for single attP and FRT landing sites have been generated (Figure 1-figure supplement 2).

The general principle we demonstrate in this study is that any genomic locus can be engineered in a scarless manner if a DNA fragment can be integrated nearby. Due to the highly conserved homologous recombination pathways, we expect this principle to be applicable to other organisms.

## Acknowledgements

We would like to thank Yikang Rong for the hs-I-SceI fly line, Al Handler for the pXLBacII-pUbDsRed-T3 plasmid, and Susan Parkhurst for the p[sChFP] plasmid. We thank Timothy J. Dahlem for his help with the design and production of the TALEN encoding plasmids. We also want to thank all past and present members in the Mann lab for insightful discussions. This study was supported by NIH Grant 5R21NS105507-02 to WBG and NIH grant R35GM118336 to RSM.

## Materials and Methods

### A. Materials

Restriction enzymes, CIP, Klenow fragment, T4 DNA polymerase and T4 DNA ligase were purchased from the New England Biolabs. Oligos were all purchased from Fisher Scientific. DH5alpha (competent cells made in house), and Stbl2 cells (Invitrogen 10268019) were the *E. coli* strains used for cloning. TALEN plasmids were designed by and purchased from University of Utah Mutation Generation and Detection Core Facility. DNA Molecular Weight Marker II, DIG-labeled (Roche 11218590910) was used as marker for all Southern blot experiments.

#### Commercial kits

AmpliScribe SP6 Transcription Kit (Epicentre AS3106).

ScriptCap m^7^G Capping System (Cellscript C-SCCE0625).

DIG High Prime DNA Labeling and Detection Starter Kit II (Roche 11585614910)

DIG Wash and Block Buffer Set (Roche 11585762001)

#### Plasmids

pBluescript II KS(+)

pUAST (Brand and Perrimon, 1993)

pUASTattB (Bischof et al., 2007)

p[sChFP] (Abreu-Blanco et al., 2012)

pUChsneo-Act (DGRC 1210) (Thummel et al., 1988)

pH-Stinger (DGRC 1018) (Barolo et al., 2000)

The MiMIC vector pMiLR-attP1-2-yellow-SA-EGFP (DGRC 1321) (Venken et al., 2011)

pXLBacII-pUbDsRed-T3 (a gift from Al Handler) (Handler and Harrell, 2001)

Addgene plasmid No. 26224

#### Flies

From Bloomington:

5905 (isogenic *w*^*1118*^)

6936 (*P{v, hs-I-CreI}; ry*^*506*^),

19139 *(w*^*1118*^; *P{w[+mC]=XP}Ubx*^*d00281*^*/TM6B, Tb*^*1*^)

36313 (*y*^*1*^, *M{RFP[3xP3.PB] GFP[E.3xP3]=vas-int.B}ZH-2A w*; Sb*^*1*^*/TM6B, Tb*^*1*^)

28877 (*lig4*)

24482 (*y*^*1*^, *M{RFP[3xP3.PB] GFP[E.3xP3]=vas-int.Dm}ZH-2A w*; M{3xP3-RFP.attP’}ZH-51C*)

33187 (*Antp*^*MI02272*^)

55598 (*MI11240*)

w-; P{v, hs-I-SceI}, Sco/CyO (a gift from Yikang Rong).

### B. Methods

#### 1. The design and optimization of the targeting vectors

3 variants of the targeting vector were designed, one for use with landing sites containing an inverted attP cassette (pTargeting-RMCE), which had from the left to the right the following elements: attB-FRT, I-CreI-mini-white-I-CreI, MCS, I-SceI-hsneo-3xP3-RFP-I-SceI, attB (Figure 1-figure supplement 1A). The other two vectors were for use with landing sites with a single attP or FRT site (pTargeting-(+) and pTargeting-(-)), which did not have the right most attB element, and differ by the orientation of the left most attB-FRT element relative to other elements in the vectors. It is worth noting that for the FRT sequence, both orientations have been defined as “positive” by different researchers, so it is important to inspect the actual FRT sequence in the landing site. In addition, the *3xP3-RFP* marker in all targeting vectors has a single loxP site (irrelevant to genome editing), which was present in the PCR template from which the *3xP3-RFP* marker was amplified.

The initial pTargeting-RMCE vector was used to generate a targeting plasmid to engineer the *Antp* locus using the *Antp*^*MI02272*^ *MiMIC* insertion as the landing site, and RMCE events were identified by the loss of the *yellow* marker, which marks the original *MiMIC* insertion. The *hs-neo* marker worked as expected, conferring G418 resistance to the RMCE flies. However, there was only very weak RFP expression in the eyes, and no *white* expression could be detected. The weak RFP expression was due to the upstream *hs-neo* element, which is expected to be transcribed through the *3xP3-RFP* marker gene because it did not have a transcription termination signal. This problem was solved by adding an SV40-polyA element downstream of *hs-neo* and upstream of *3xP3-RFP*. This SV40-polyA element was included in all subsequent targeting vectors.

The lack of *mini-white* expression is most likely due to it being silenced in the *Antp* locus. The main evidence supporting this explanation is that all *mini-white* insertions that are expressed within the *Antp* locus are flanked by insulator elements. A similar observation was also seen in the *Bithorax Complex*, where the *Hox* gene *Ubx* resides. Within the *Bithorax Complex*, all *mini-white* marked transposons also have insulated *mini-white*, while immediately outside of the *Bithorax complex*, both insulated and non-insulated *mini-white* genes are expressed. Consistently, the non-insulated *mini-white* marker is silenced when inserted into the majority of genomic loci (Handler and Harrell, 1999, Horn et al., 2000). Insulated targeting vectors were thus generated, in which 4 *gypsy* insulators were added to each version of the vectors, 2 flanking *mini-white*, and 2 flanking *hsneo-3xP3-RFP*. There are repetitive sequences in the gypsy insulators, and two insulators near each other would make the plasmids unstable. Therefore, a ∼2 kb spacer (from the *Drosophila yellow* gene) was inserted into the middle of the MCS in all insulated targeting vectors, to separate the right *mini-white* insulator from the left *hsneo-3xP3-RFP* insulator (Figure S1A). Because of the presence of multiple insulators, Stbl2 *E. coli* cells, which increases the stability of plasmids with repetitive sequences, must be used to manipulate the insulated targeting vectors. All preps of targeting plasmids derived from insulated targeting vectors should be verified by restriction digestion verification before being used in injection, as plasmid rearrangement happens more frequently during the growing of large volume *E. coli* cultures.

The details of targeting vector cloning are in Supplementary file 1, and the sequences of all targeting vectors are in Supplementary file 3.

#### 2. The generation of the *Ubx* landing site line

To generate a custom landing site in the Ubx locus between the ATG start codon and the W-motif codons, a pair of TALENs were designed. To avoid potential issues caused by natural polymorphisms, this *Ubx* region of the *lig4* strain (Bloomington #28877), which would be used in TALEN-mediated genome targeting, was PCR amplified and sequenced. The exact sequence in the *lig4* strain was sent to the University of Utah Mutation Generation and Detection Core Facility for identification of optimal TALEN target sites, and the most promising pair of TALENs were then purchased. The TALEN target sequence is: TGCCCGTTAGACCCTCCGCCT-gcaccccagattcccg-AGTGGGCGGCTATTTGGA, in which the upper-case letters show the TALEN binding sites, and the lower-case letters indicate the spacer between the two binding sties.

The TALEN plasmids were linearized by restriction digestion and gel purified, and were used as templates for *in vitro* transcription using the AmpliScribe SP6 Transcription Kit (Epicentre AS3106). The mRNAs were then capped in a subsequent reaction using the ScriptCap m^7^G Capping System (Cellscript C-SCCE0625).

A vector, *pCassette-ubiDsRed*, was generated, which has an *ubiDsRed* marked inverted attP cassette flanked by two different multiple cloning sites (MCS) for inserting homologous arms. Ubx-N-L and Ubx-N-R homologous arms were cloned into these two MCS sites to generate the *pCassette-Ubx-N* donor plasmid. A mixture of this donor plasmid (final concentration 500 ng/ul) and the two capped TALEN mRNAs (final concentration 400 ng/ul each) was injected into the blastoderm of *lig4* embryos (injection done by BestGene Inc.), and the desired homologous events were identified by strong ubiquitous DsRed expression in the F1 generation. Positive individuals were used to generate stocks and several independent stocks were verified by Southern blot and sequencing. One fully verified line was used as *Ubx* landing site line.

The sequences of the TALEN plasmids are in Supplementary file 3, and the detailed cloning steps for the landing site donor plasmid are in Supplementary file 1.

#### 3. Building suitable homing nuclease-expressing fly strains

Standard fly genetics was used to mobilize P element to obtain *hs-I-SceI(X)* and *hs-I-CreI(II)*. Because the hs-I-SceI transgene was marked with *vermillion (v)*, an attempt was made to generate the line *v*^*1*^; *P{v, hs-I-SceI}, Sco/CyO* for P element mobilization from the strain *w-; P{v, hs-I-SceI}, Sco/CyO* (a gift from Yikang Rong). However, *v*^*1*^*/(FM7C); P{v, hs-I-SceI}, Sco/CyO* females were sterile, so instead, the line *v*^*1*^; *Pin, P{v, hs-I-SceI}/CyO* was generated, and was used as the starting line to jump P{v, hs-I-SceI} from chromosome II to X chromosome. The *P{v, hs-I-CreI}* P element was jumped to chromosome II from the X chromosome, using *v*^*1*^, *P{v, hs-I-CreI}; ry*^*506*^ (Bloomington #6936) as the starting line.

X chromosome with the genotype *v*^*1*^, *P{v, hs-I-SceI}, P{v, hs-I-CreI}*, and chromosome II with the genotype *Pin, P{v, hs-I-SceI}, P{v, hs-I-CreI}* were then generated by recombination. *v+* recombinants were screened for the presence of both *hs-I-SceI* and *hs-I-CreI* transgenes by PCR using primer pairs *hs-I-SceI-5’ + hs-I-SceI-3’*, and *hs-I-CreI-5’ + hs-I-CreI-3’*, respectively. Finally, appropriate balancers were added by crossing.

All primer sequences are in Supplementary file 2.

#### 4. Cloning of the integrated fragments

For *Antp* and *Ubx* targeting, the integrated fragment was assembled from 3 sub-fragments, and the ∼2 kb middle sub-fragment contained the loci to be mutated. The 3 sub-fragments were PCR amplified from genomic DNA and cloned into the pBluescript vector. Both the PCR products and the cloned fragments were fully sequenced to ensure no PCR-introduced mutations in the cloned fragments. The desired mutations were then introduced to the middle sub-fragment by standard procedures. Next, a 3-fragment ligation was performed to assemble the complete integrated fragment in pBluescript. The assembled integrated fragment was then cloned into the targeting vector *pTargeting-RMCE-insulated*. For *Gr28b* deletion, a 2kb left arm and a 2.3kb right arm flanking the desired deletion were PCR amplified from genomic DNA, digested with restriction enzymes, and ligated into pBluescript in a 3-fragment ligation reaction. The 4.3 kb integrated fragment was then cloned into the targeting vector *pTargeting-RMCE-insulated*.

All detailed cloning steps are in Supplementary file 1.

#### 5. The identification and verification of RMCE alleles

All the landing site lines were verified before being used in injections. The *Ubx* landing site line was generated in this study, and was fully verified by Southern blot analysis and sequencing. For *Antp* and *Gr28b* targeting, *MiMIC* insertions were used as landing sites, and the presence of the desired *MiMIC* insertions was verified by PCR. Clean genetic sublines, which removed a linked lethal mutation, were derived from single *Antp*^*MI02272*^ chromosomes, and one was selected for all subsequent injections.

Initially, *vas-int(X); MiMIC* stocks were generated and tested for injection, but the injected embryos suffered high fatality rates. Improved survival was obtained from injecting the F1 embryos of the crosses between the *vas-int(X)* females and landing site containing males. All injections were done by BestGene Inc. G0 adults from the injected embryos were individually crossed to suitable balancer stocks, and the F1 flies were screened for RMCE events. The RMCE alleles were identified by the presence of the *mini-white* marker, and the presence of *3xP3-RFP* and the loss of the original landing site marker were then confirmed for all *white+* individuals. RMCE stocks were then established from individual flies with the correct marker patterns. The orientations of the RMCE lines were determined by PCR.

At first, RMCE alleles were verified by Southern blotting before they were used for resolution. Later, a more efficient procedure was used: several RMCE alleles with the correct marker patterns were subjected to resolution without Southern blot verification, and fewer individual cross IIs from each RMCE allele were set up. After getting all final mutant alleles, Southern blot analyses of the RMCE alleles were performed alongside with selected final mutant alleles. This arrangement also enables more independent mutant alleles to be obtained.

During injections, 2 classes of abnormal recombination events were observed. 1. Some transformants had both *mini-white* and *3xP3-RFP*, but the original landing site marker (*yellow* or *ubiDsRed*) remained present. These events probably resulted from site-specific recombination between a single pair of attP and attB sites, whereas the other recombination events did not happen. Or maybe two different plasmids were integrated into the genome, each via one site-specific recombination event. 2. As mentioned in the Results section, some transformants lost the landing site marker, but only *mini-white* was present, and no *3xP3-RFP* was observed. This class was most likely because of spontaneous resolution of the right end during phiC31 integrase mediated RMCE, in which dsDNA breaks were introduced within the attP and attB sites, and could have triggered homologous recombination. The RMCE transformants were usually selected by the presence of *mini-white*, and the presence of *3xP3-RFP* and the absence of the landing site marker were confirmed later. Therefore, it is reasonable to expect that *3xP3-RFP+, white-, yellow-(ubiDsRed-)* transformants also existed, but they were unidentified. Spontaneous resolution of both ends during RMCE might also happen at low frequency.

The primer sequences for verifying *MiMIC* and RMCE alleles are in Supplementary file 2.

#### 6. Resolving the RMCE alleles to generate the final mutant alleles, and the definition of productive cross IIs

The crosses to resolve RMCE alleles of different chromosomes are shown in Figure 3, Figure 4A and Figure 4-figure supplement 1. All crosses were performed at 25°C. The following describes details of the resolution steps for chromosome II or III targets. If the target is on the X chromosome, individual females must be used in Cross IIs and Cross IIIs, and some details should be adjusted accordingly.

For Cross I, several vials of crosses were set up, and the flies were allowed to accommodate for a few days. The adults were then allowed to lay embryos for 72 hours before being transferred to new vials, and the embryo/larvae in the old vials were heat shocked at 37°C. If I-SceI was the only homing nuclease expressed, 1-hour heat shock was performed. A 20-minute heat shock was performed if I-CreI was involved, either with or without I-SceI (Note: in the sequential resolution reported here, a 40-minute heat shock was performed to induce I-CreI expression, but later results showed that a 20-minute heat shock might give better efficiency). A second 72-hour collection and heat shock might be performed if necessary. When the heat shocked individuals reach adult stage, males of the desired genotype were individually crossed to a balancer line in Cross II. The progeny of Cross IIs was screened once every 2 to 3 days for males that lost the desired marker(s). For simultaneous resolution, white-eyed males were first identified, and the *3xP3-RFP* marker was then inspected under a fluorescent scope. Once male progeny that lost the desired marker(s) was identified from a Cross II, this particular Cross II was not screened further. To ensure all final alleles were independent, for each Cross II, only one Cross III was set up. If the selected individual male used in a Cross III turned out to be sterile, no extra Cross IIIs were set up for the corresponding Cross II, even if that Cross II might have produced more males that lost the desired marker(s).

For the purpose of easy scoring and comparison, a productive Cross II was defined as an individual Cross II that eventually generated a final stock. Occasionally, the selected single male from a Cross II was sterile, and this particular Cross II would be scored as non-productive. In some cases, the final stock from a Cross II might not be a correctly resolved allele (for example, it might be a right marker deletion event), but such a Cross II would be scored as productive according to the above definition.

#### 7. Southern blot analysis

Southern blots were performed using the DIG High Prime DNA Labeling and Detection Starter Kit II (Roche 11585614910) and the DIG Wash and Block Buffer Set (Roche 11585762001), according to manufacturer’s instructions. DNA Molecular Weight Marker II, DIG-labeled (Roche 11218590910) was used as marker. In general, two probes were needed to verify the selected alleles. After hybridizing with the first probe, the blot was stripped and re-hybridized with the second probe according to manufacturer’s instructions. For *Antp* and *Ubx* targeting, the left and right sub-fragments in the integrated fragment (see above) were used to generate DIG labeled 5’ and 3’ Southern blot probes. For *Gr28b* deletion, the left and right arms (see above) were used as templates to generate the probes.

#### 8. Sequencing of the mutant alleles

For all selected final mutant alleles, the genomic region corresponding to the integrated fragment in the targeting plasmid plus short (100-200bp or so) flanking regions was completely sequenced. For homozygous lethal alleles, embryos were collected overnight at 25°C from the balanced stock, and were further aged at 25°C for at least 30 hours. 6 unhatched embryos were randomly selected and single embryo genomic DNA extraction was performed. A fragment covering the regions with desired mutation(s) was PCR amplified and sequenced to genotype the selected embryos. Homozygous mutant embryos were identified and their genomic DNA samples were used as PCR templates. For homozygous viable alleles, homozygotes were used to extract genomic DNA. The region to be sequenced was divided into 2-3kb fragments with small overlaps. These fragments were PCR amplified with Phusion DNA polymerase, and gel purified before sequencing with sequence specific primers. Gel purification was necessary to obtain high quality sequencing results, especially if the genomic DNA was from single embryos.

In all the targeting cases reported in this study, there are natural polymorphisms between the landing site line and the line from which the donor fragment was PCR amplified. The pattern of polymorphisms in the resolved lines generally showed the expected pattern: the landing site-proximal regions often had the polymorphisms from the integrated fragments, while the landing site-distal regions usually had the polymorphisms from the original landing site line.

#### 9. Calculating the fraction of the fly genome accessible by our technique

In this study, we calculated the fraction of the fly genome that can be accessed by our technique from a mapped *MiMIC* insertion, assuming 5 kb near a landing site can be reached without difficulty. The *MiMIC* insertions were selected for calculation because these are among the most efficient landing sites and there are well-documented *MiMIC* mapping data at the base pair resolution.

The *MiMIC* mapping results were downloaded from the URL: http://flypush.imgen.bcm.tmc.edu/pscreen/downloads.html. The original file contains the base pair positions of 7441 *MiMIC* insertions, all of which are in euchromatic regions. 9 insertions with incomplete mapping information were dropped, leaving 7432 insertions with complete mapping information. A bed file containing 10 kb genomic intervals centered at each of these 7432 *MiMIC* insertions was then generated. Next, the “MergeBED” function in bedtools (performed on usegalaxy.org) was used to generate a new bed file that contains 4154 non-redundant genomic intervals covering all sequences equal to or less than 5kb from one of the 7432 *MiMIC* insertions. The length (in base pair) of each of these 4154 genomic intervals was then calculated in Microsoft Excel. Finally, the total length of all 4154 intervals was calculated to be 57,528,100 bp, which is roughly half of the fly genome that is euchromatic (117 Mb) (Hoskins et al., 2015).

Since the mapped *MiMIC* lines represent only a subset of all available landing sites, and the estimate that 5 kb flanking a landing site can be engineered is a conservative one, the actual fraction of accessible fly genome is expected to be significantly larger than 50%.

## Supplementary figures

**Figure 1-figure supplement 1.**
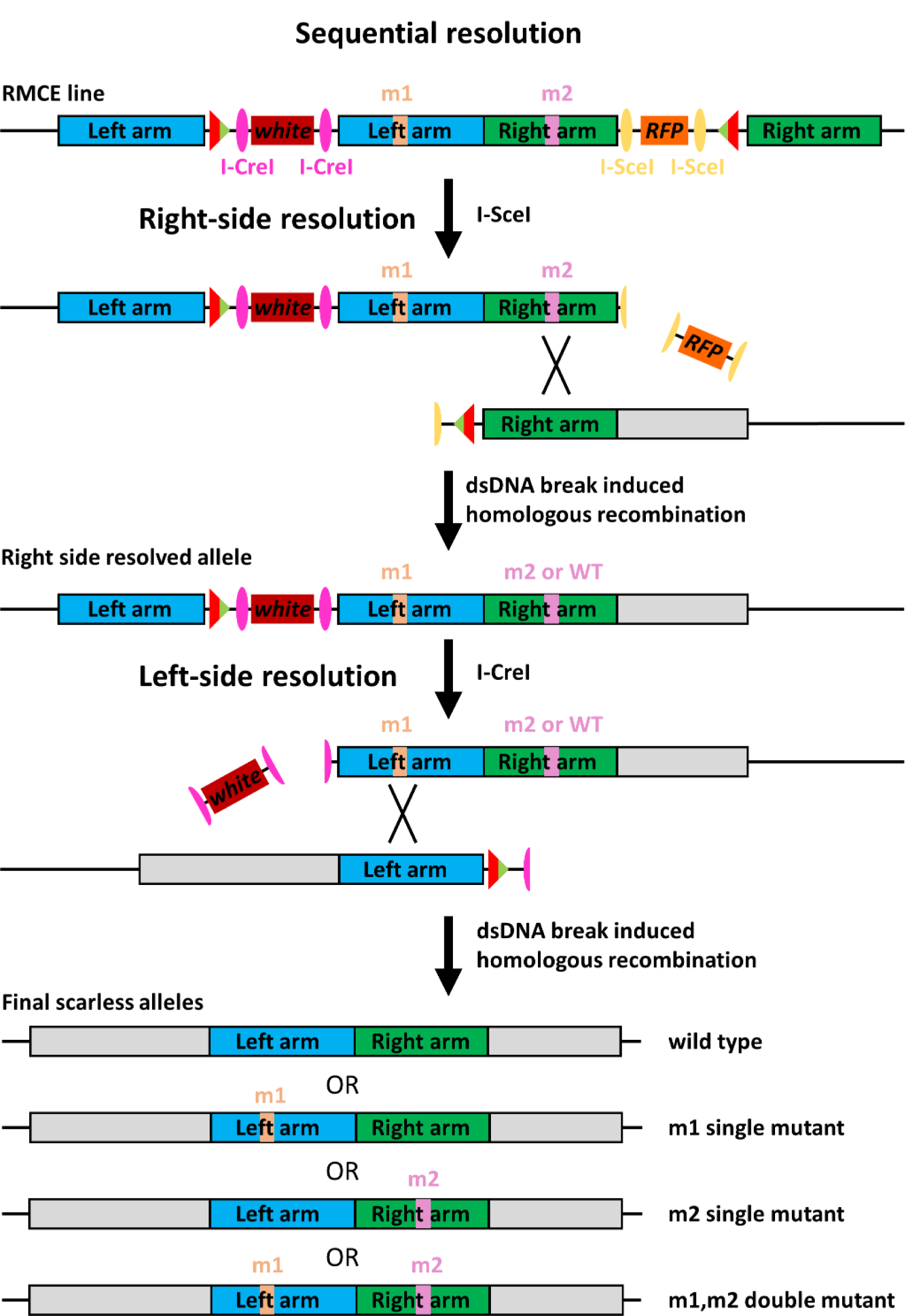
Sequential resolution of the RMCE line. Schematics showing sequential resolution of the RMCE allele. The right side is first resolved by I-SceI expression, followed by left-side resolution by the expression of I-CreI.

**Figure 1-figure supplement 2.**
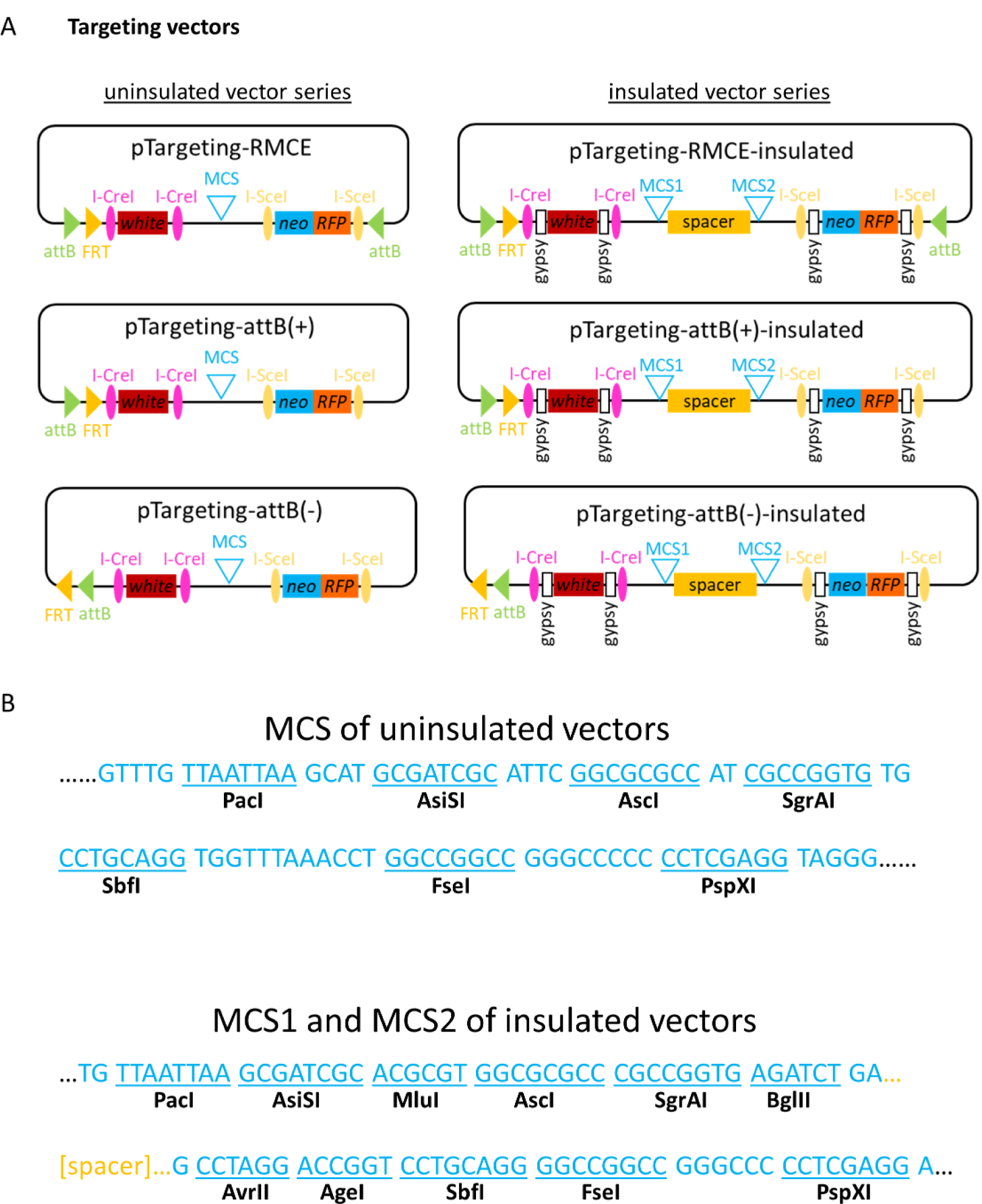
Targeting vectors. **A.** Maps of targeting vectors. The RMCE vectors are for landing sites containing inverted attP cassette, and the attB vectors are for landing sites with a single attB (or FRT) site. The (+) and (-) versions differ in the orientations of the attB and FRT sites relative to the rest of the vector. The uninsulated series are suitable for loci where *mini-white* is known to be expressed, and the insulated series should be used in loci where *mini-white* is or might be silenced. One way to determine if *mini-white* is silenced in the locus of interest is to examine existing *mini-white* marked transposon insertions near the locus of interest. If all such transposons contain insulated *mini-white, mini-white* is likely to be silenced near this particular locus in the genome. See Materials and Methods for more details on the silencing of *mini-white*. **B.** The multiple cloning site regions of the targeting vectors. Only unique restriction sites are indicated. For insulated vectors, the 2 kb spacer separates the two flanking insulators and reduces plasmid instability during cloning. One site from MCS1 and one from MCS2 should be selected when using the insulated vectors to remove the spacer.

**Figure 2-figure supplement 1.**
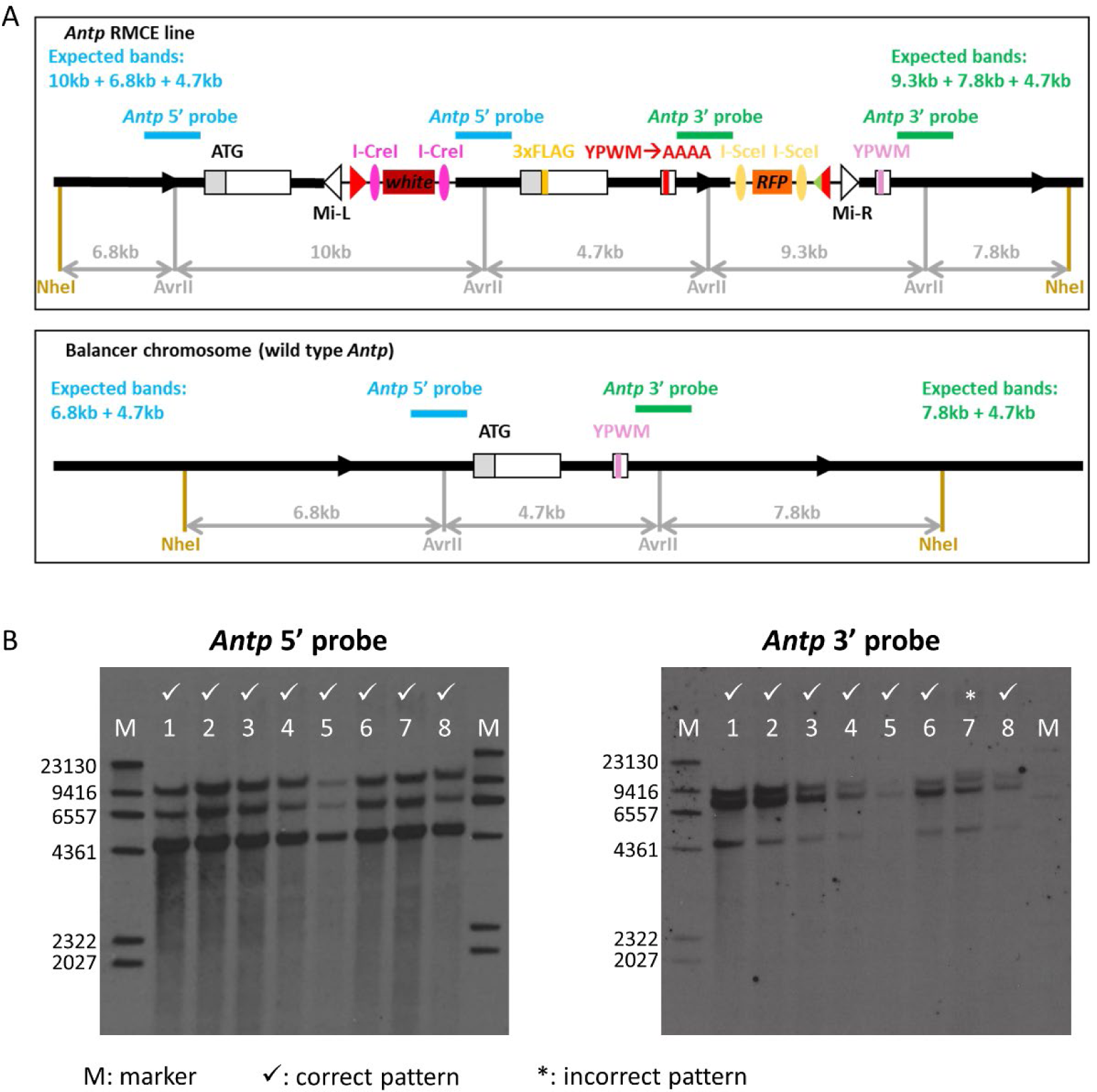
Southern blot verification of multiple independent *Antp* RMCE alleles. **A.** Restriction maps of the *Antp* RMCE allele and the wild type *Antp* allele (the balancer chromosome). The positions of relevant restriction sites and the sizes of all relevant restriction fragments are shown. The regions used as 5’ and 3’ Southern blot probes are indicated with blue and green bars. The expected Southern blot patterns for each probe are also shown. The schematics are not drawn to scale. **B.** Southern blot results for 8 independent *Antp* RMCE alleles. The *Antp* RMCE alleles are homozygous lethal and are balanced with a balancer chromosome, which contributes to the observed Southern blot patterns. Sample 7 has an additional band above the 9.3 kb band when blotted with the *Antp* 3’ probe, indicating it might have additional rearrangement(s).

**Figure 3-figure supplement 1.**
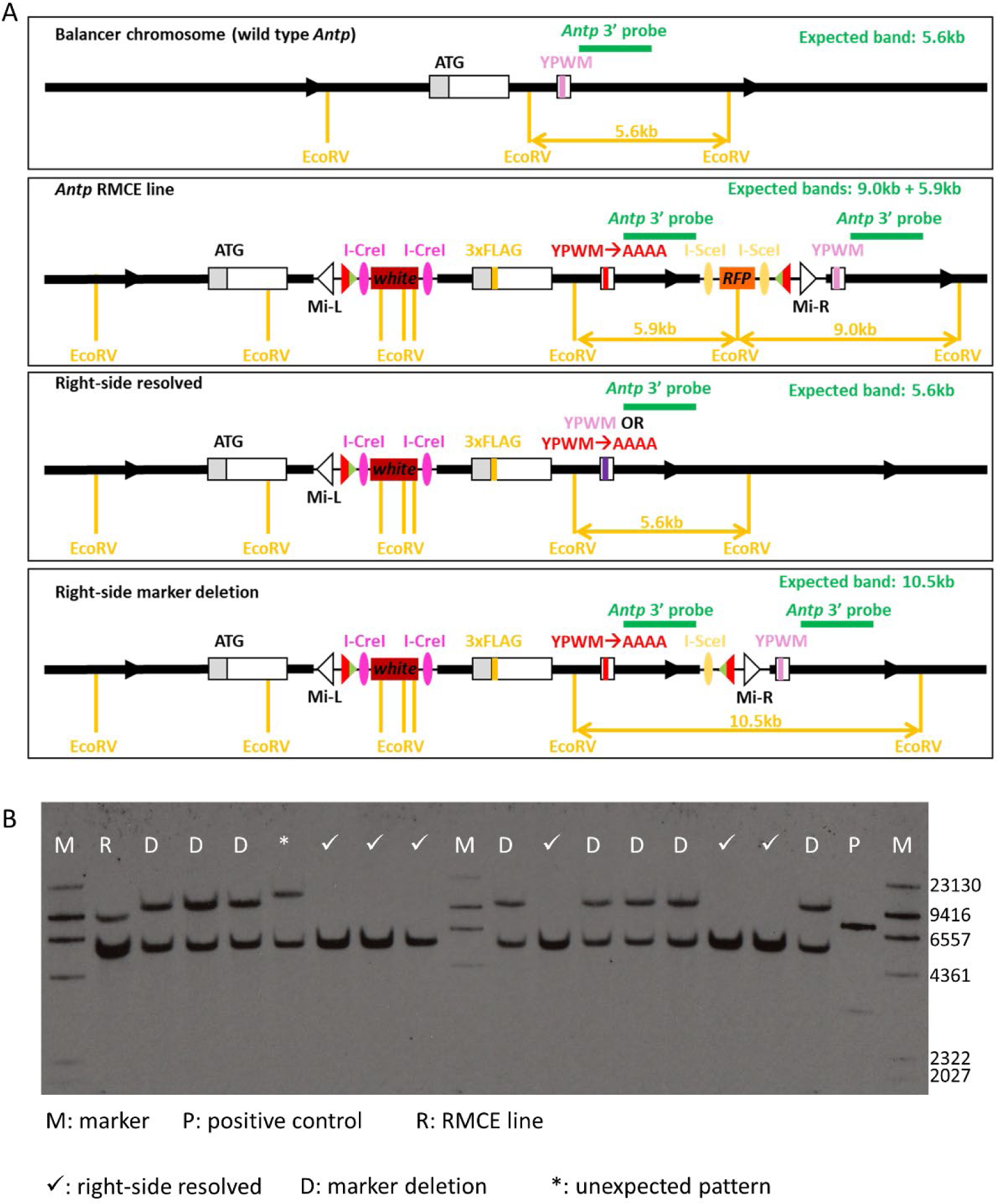
Southern blot analysis of alleles from I-SceI mediated right-side resolution. **A.** Restriction maps of various genotypes. The positions of relevant restriction sites and the lengths of relevant restriction fragments are indicated. The green bar indicates the region used as the *Antp* 3’ probe, and the expected Southern blot pattern for each genotype is also shown. The schematics are not drawn to scale. **B.** Southern blot analysis of a subset of selected alleles. All alleles are homozygous lethal and are balanced with a balancer chromosome, which gives a 5.6 kb band. The positive control is restriction digested plasmid with a fragment from *Antp*.

**Figure 3-figure supplement 2.**
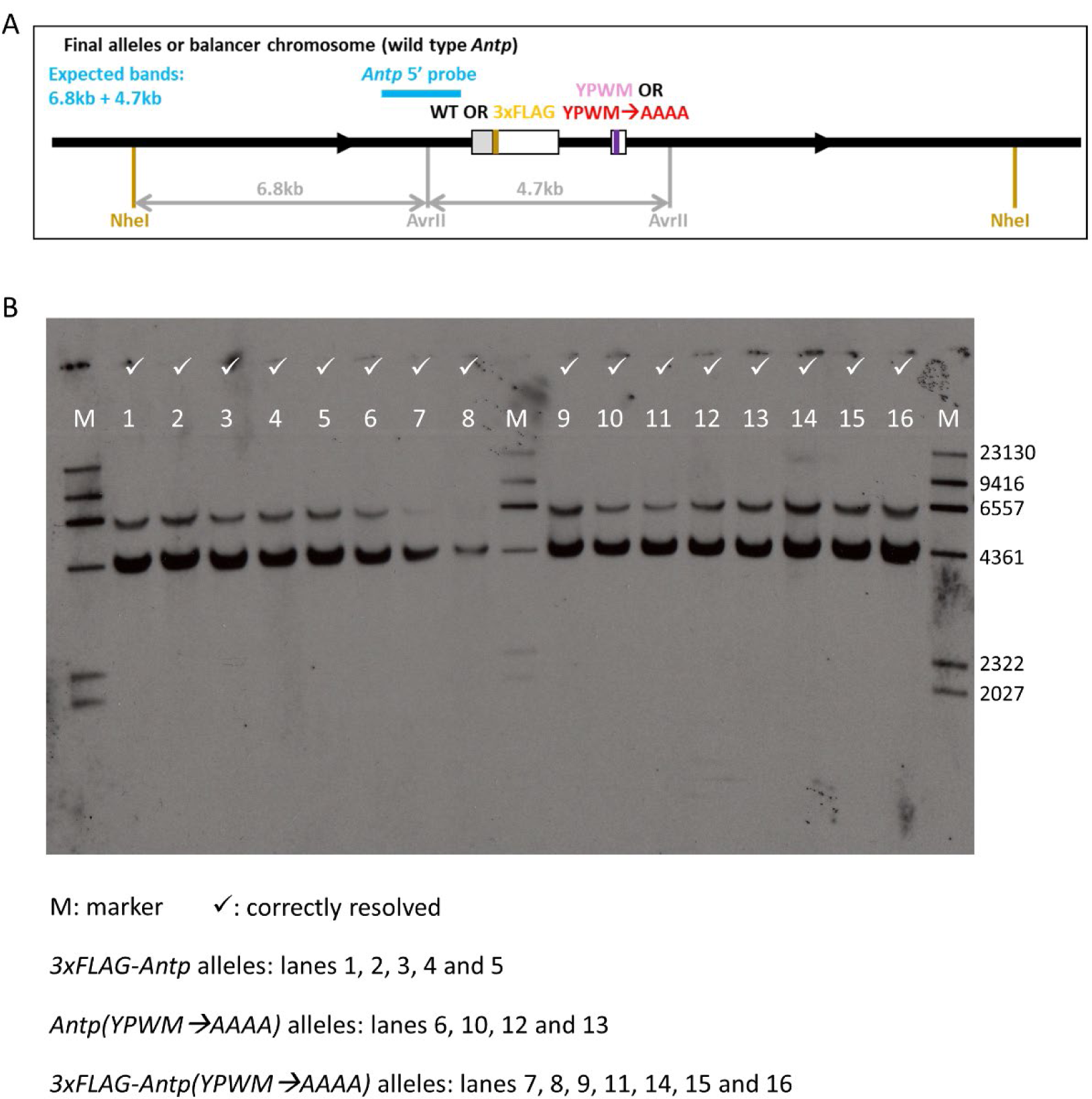
Southern blot verification of final *Antp* alleles from sequential resolution. **A.** Restriction map of the correct final *Antp* alleles. The positions of the relevant restriction sites are indicated, as well as the lengths of relevant restriction fragments. The blue bar shows the region used as the *Antp* 5’ Southern blot probe. The expected Southern blot pattern is also indicated. This schematic is not drawn to scale. **B.** Southern blot results for 16 selected final alleles. The genotype of each alleles is shown below the blot and all alleles are balanced or are segregating a balancer chromosome. All alleles on this blot show the expected pattern. Lanes 7 and 8 both had the correct patterns; the weak large molecular weight bands were confirmed by prolonged exposure (not shown).

**Figure 4-figure supplement 1.**
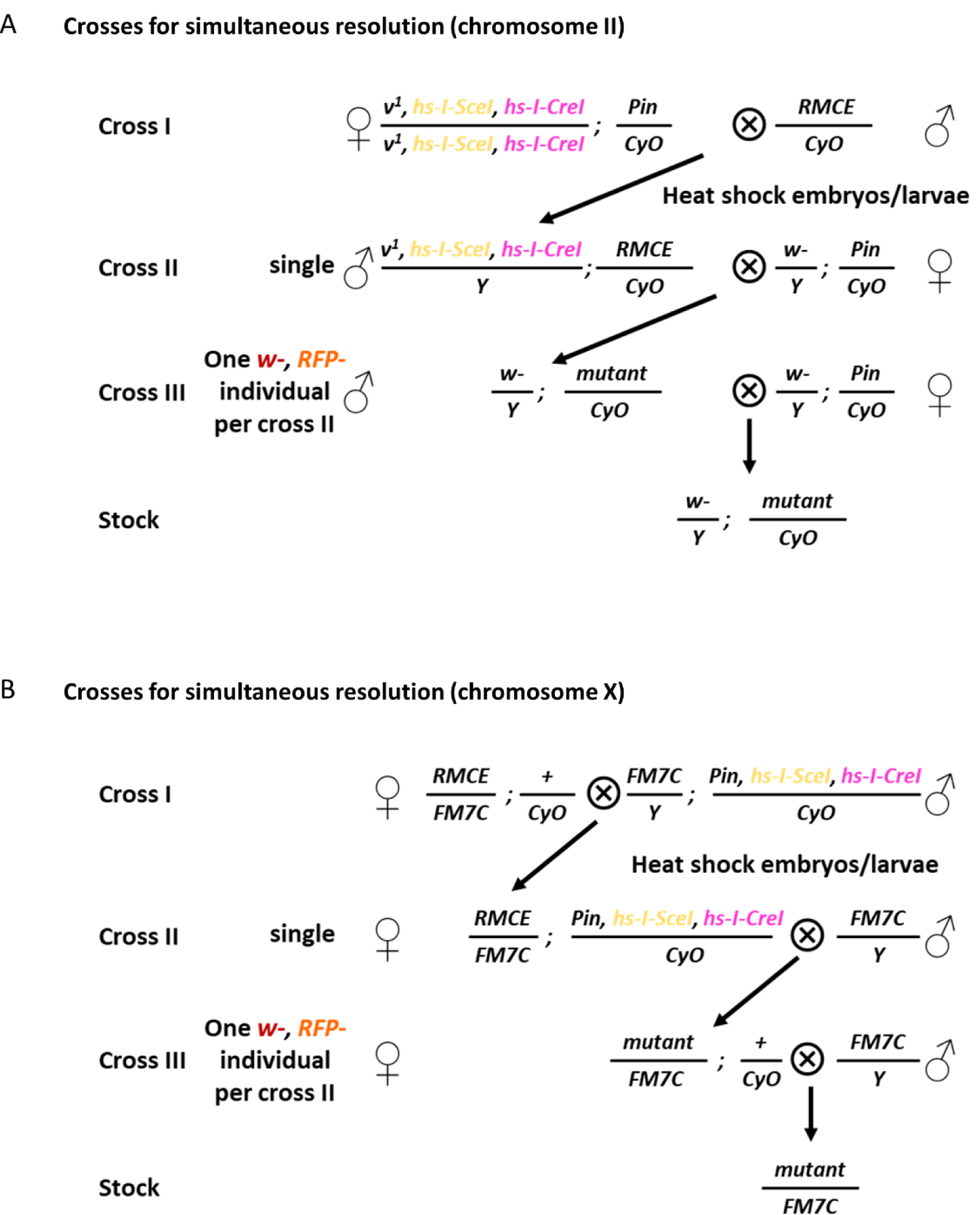
Crosses for the simultaneous resolution of second and X chromosome RMCE alleles. **A.** Crosses for simultaneously resolving RMCE alleles on chromosome II. **B.** Crosses for the simultaneous resolution of X chromosome RMCE alleles.

**Figure 4-figure supplement 2.**
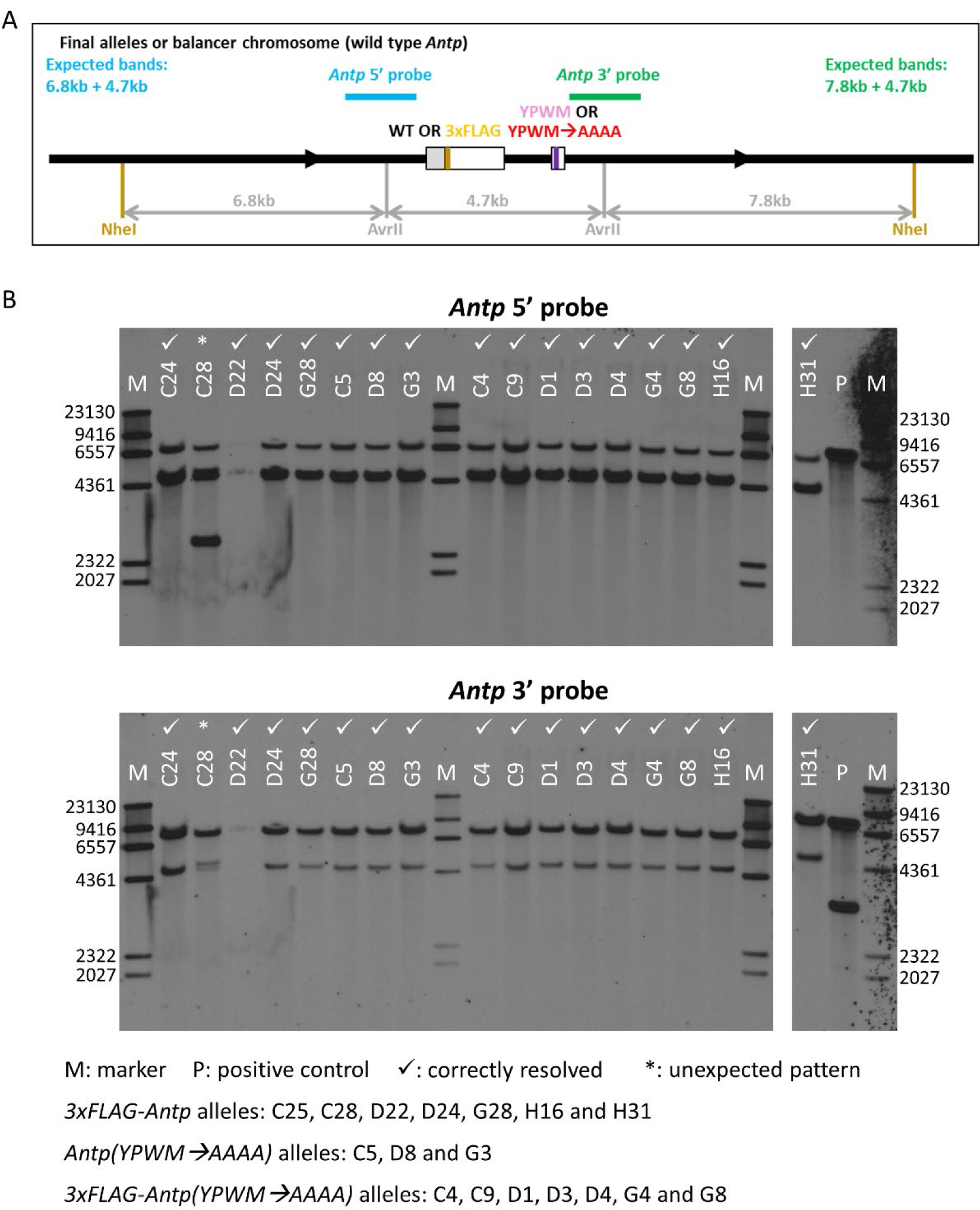
Southern blot verification of final alleles from the simultaneous resolution of *3xFLAG-Antp* RMCE alleles. **A.** Restriction map of the correct final alleles. The positions of relevant restriction sites, and the lengths of relevant restriction fragments are shown. The blue and green bars indicate the regions used as *Antp* 5’ and *Antp* 3’ Southern blot probes. The expected Southern blot patterns from each probe are also shown. This schematic is not drawn to scale. **B.** Southern blot results of selected final alleles. The genotype of each allele is shown below the blots, and all alleles are balanced with or are segregating a balancer chromosome. The letter in the name of a final allele indicates the original RMCE line from which this allele was derived. Other than C28, all alleles show the correct pattern. D22 had weak signal, but its correct pattern was confirmed by prolonged exposure (not shown). The positive control is restriction digested plasmid with a fragment from *Antp*.

**Figure 5-figure supplement 1.**
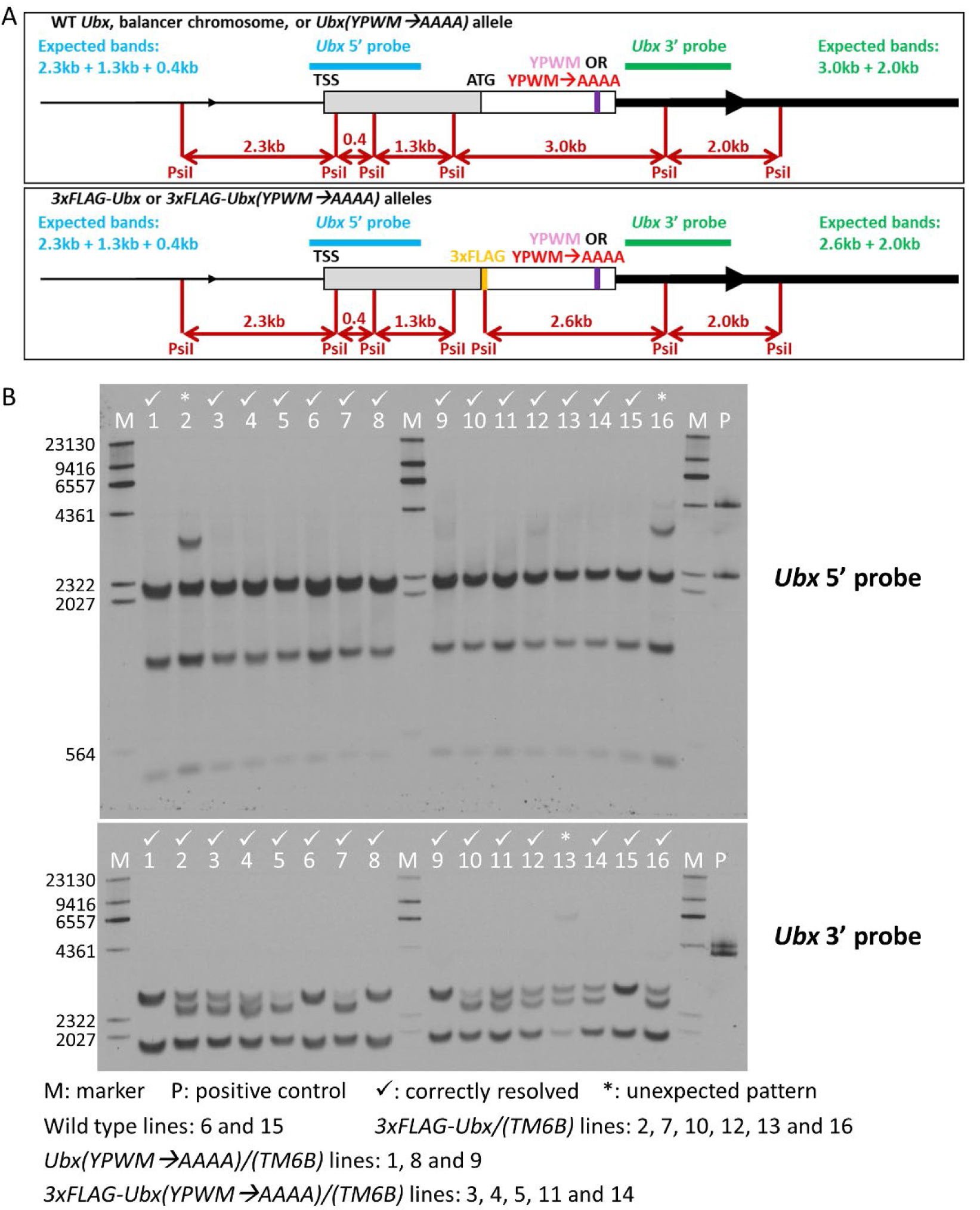
Southern blot verification of selected final *Ubx* alleles. **A.** Restriction maps of final *Ubx* alleles. The positions of relevant restriction sites are shown, as well as the sizes of the relevant restriction fragments. The blue and green bars indicate regions used as the *Ubx* 5’ and *Ubx* 3’ probes. The expected Southern blot patterns for each probe are shown. The schematics are not drawn to scale. **B.** Southern blot results of selected final *Ubx* alleles. The genotype of each sample is shown below the blots. All lines might be segregating a balancer chromosome. Lanes 2, 13 and 16 each showed an extra band, indicating additional rearrangement(s). The positive control is restriction digested plasmid with a fragment from *Ubx*.

**Figure 6-figure supplement 1.**
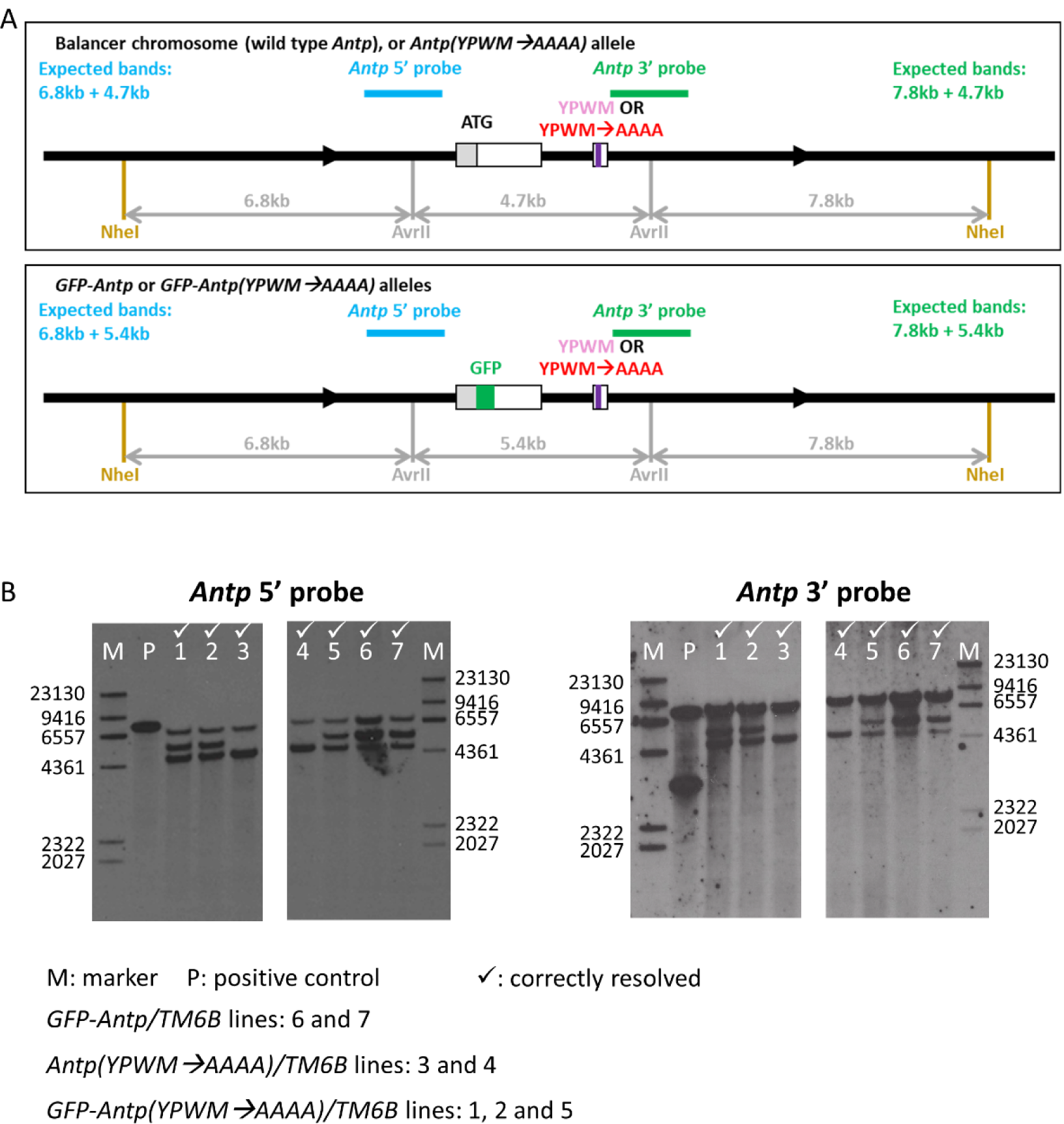
Southern blot verification of selected *GFP-Antp* targeting final alleles. **A.** Restriction maps of various genotypes. The positions of the relevant restriction sites are indicated, and the lengths of the relevant restriction fragments are also shown. The blue and green bars show the regions used as *Antp* 5’ and *Antp* 3’ Southern blot probes. The expected patterns from each probe are indicated. These schematics are not drawn to scale. **B.** Southern blot results of selected final *GFP-Antp* targeting alleles. The genotype of each sample is indicated below the blots. All of these alleles give correct patterns. The positive control is restriction digested plasmid with a fragment from *Antp*.

**Figure 7-figure supplement 1.**
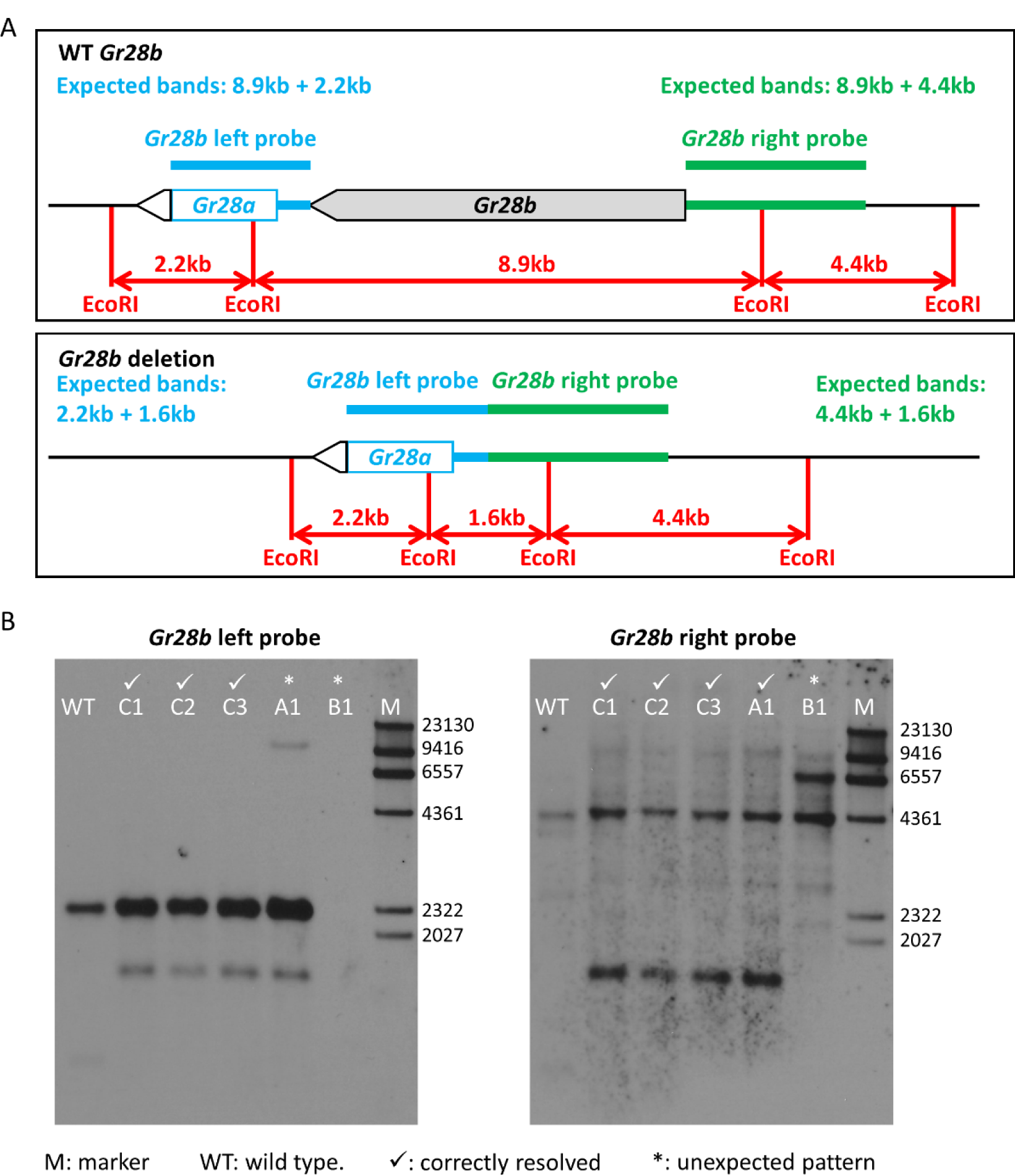
Southern blot verification of selected *Gr28b* deletion alleles. **A.** Restriction maps of the wild type *Gr28b* allele and the *Gr28b* deletion allele. The positions of relevant restriction sites, as well as the lengths of relevant restriction fragments, are shown. The blue and green bars indicate the regions used as the *Gr28b* left and *Gr28b* right Southern blot probes. The expected patterns from each probe are also shown. Not drawn to scale. **B.** Southern blot results of 5 selected final *Gr28b* deletion alleles. Genomic DNA was extracted from homozygous flies for all samples. For the wild type control, the 8.9 kb band is probably too weak to be visible. Alleles C1, C2 and C3 are correct. Allele A1 has an extra band when probed with the *Gr28b* left probe. Allele B1 does not show any signal when probed with the *Gr28b* left probe, indicating sequences homologous to this probe are absent. The additional weak bands visible in the blot probed with *Gr28b* right probe may be due to repetitive sequences in the probe.

